# Pharmacologically targeting a novel pathway of sodium iodide symporter trafficking to enhance radioiodine uptake

**DOI:** 10.1101/622241

**Authors:** Alice Fletcher, Martin L. Read, Caitlin E.M. Thornton, Dean P. Larner, Vikki L. Poole, Katie Brookes, Hannah R. Nieto, Mohammed Alshahrani, Rebecca J. Thompson, Gareth G. Lavery, Moray J. Campbell, Kristien Boelaert, Andrew S. Turnell, Vicki E. Smith, Christopher J. McCabe

## Abstract

Radioiodine treatment fails ≥25% of patients with thyroid cancer and has been proposed as a potential treatment for breast cancer. Cellular iodide uptake is governed by the sodium iodide symporter (NIS), which is frequently mislocalized in thyroid and breast tumours. However, the trafficking of NIS to the plasma membrane (PM) is ill-defined. Through mass spectrometry, co-immunoprecipitation, cell surface biotinylation and proximity ligation assays we identify two proteins which control NIS subcellular trafficking: ADP-ribosylation factor 4 (ARF4) and valosin-containing protein (VCP). HiLo microscopy revealed ARF4 enhanced NIS trafficking in co-incident PM vesicles, governed by a C-terminal VXPX motif, whilst papillary thyroid cancers (PTC) demonstrate repressed ARF4 expression. In contrast, VCP, the central protein in ER-associated degradation, specifically bound NIS and decreased its PM localization. Five chemically distinct allosteric VCP inhibitors all overcame VCP-mediated repression of NIS function. In mice, two re-purposed FDA-approved VCP inhibitors significantly enhanced radioiodine uptake into thyrocytes, whilst human primary thyrocytes showed similar increases. Critically, PTC patients with high tumoural VCP expression who received radioiodine had strikingly worse disease-free survival. These studies now delineate the mechanisms of NIS trafficking, and for the first time open the therapeutic possibility of systemically enhancing radioiodine uptake in patients via FDA-approved drugs.

**One Sentence Summary:** Novel NIS interactors ARF4 and VCP alter NIS trafficking *in vitro*, and FDA-approved VCP inhibitors can significantly enhance radioiodine uptake.

## INTRODUCTION

For over 75 years radioiodine treatment has been the central post-surgical therapy for patients with differentiated thyroid cancer (DTC). However, at least a quarter of DTC patients do not uptake sufficient radioiodine (^131^I) for effective ablation (*1, 2*), and this remains an urgent problem in metastatic disease. There are essentially two cohorts of thyroid cancer patients: those who respond to radioiodine and have an excellent prognosis, and those who do not respond, and for whom outcome is dire (*3*). However, no substantial changes have been made to the way radioiodine is administered therapeutically, which might improve outcome.

Troublingly, DTC is now the most rapidly increasing cancer in the UK and US, with 300,000 new cases reported worldwide per annum, and more than 40,000 people whom die from thyroid cancer annually (*4*). Mechanisms which influence treatment success in radioiodine-resistant thyroid cancers have been described in elegant but fragmented investigations. Whilst undoubted clinical progress has been made (*5-7*) the underlying causes of treatment failure are still not fully understood, and the tumoural processes which underpin radioiodine success in individual patients need to be better delineated.

The sodium iodide symporter (NIS) is the sole human transporter responsible for iodide uptake (*8*), exploitation of which represents the first – and most specifically targeted – internal radiation therapy in existence. High-energy β-emitting ^131^I is utilized to destroy remaining thyroid cells post-surgery, and target metastases. Decreased levels of NIS expression and/or diminished targeting of NIS to the plasma membrane (PM) of thyroid cancer cells represent the principal mechanisms of radioiodine-refractory disease (*9-11*). Numerous studies have addressed the common pathways of NIS regulation *in vitro* and *in vivo* (*12-17*). Other studies have addressed the key transcriptional and epigenetic alterations which silence thyroid-specific genes including NIS (*15, 18-21*). Approaches to improve the treatment of thyroid cancer have been undertaken in clinical trials including treatment with retinoids (*22*), PPARγ agonists (*23, 24*), MAPK pathway/BRAF inhibitors (*5, 6, 25*), multi-targeted kinase inhibitors (*26*) and HDAC inhibitors (*27*). Multiple biologically-targeted drugs have been evaluated in phase I, II and III trials, with several agents including sorafenib (*6*), lenvatinib (*28*) and dabrafenib (*7*) showing promising responses and/or disease stabilisation. However, issues of toxicity and drug resistance remain.

To actively transport iodide for thyroid hormone biosynthesis and radioiodine treatment, NIS must be present in the basolateral PM of thyroid follicular cells. However, relatively little is known about the mechanisms that govern NIS trafficking. TSH induces iodide uptake through upregulation of NIS expression and modulation of its subcellular localisation (*29-31*). Yet, many thyroid cancers demonstrate reduced NIS activity through diminished PM retention (*32-34*). BRAF-mutant tumours (60-70% of thyroid cancers) are more likely to be resistant to radioiodine, partly due to decreased NIS expression (*13, 35*), but also due to impaired PM targeting (*11, 14*), through mechanisms which remain ill-defined. Currently, PTTG1-binding factor (PBF) is the only protein shown to bind NIS and modulate its subcellular localisation (*36*).

Early studies identified that breast tumours can uptake radioiodine (*37, 38*), and subsequent studies confirmed functional NIS expression in up to ∼80% of breast cancers (*39-41*). However, NIS is rarely localized to the PM in breast cancers, and currently of little clinical utility (*40-43*). We hypothesized that undefined proteins interact with NIS and regulate its trafficking to, or retention at, the PM. We therefore performed mass spectrometry (MS/MS) and appraised a series of protein candidates in thyroid and breast cancer models. We report two new proteins – ADP-ribosylation factor 4 (ARF4) and valosin-containing protein (VCP) – which specifically bind NIS and directly regulate its function, providing new hope that NIS activity may be stimulated in patients who are currently radioiodine-refractory.

## RESULTS

### Identification and manipulation of NIS interactors

We performed MS/MS in MDA-MB-231 cells with lentivirally-expressed NIS (NIS+ve) to identify interactors in whole cell and PM extracts. NIS and its protein interactors were subjected to in-gel tryptic digest and resulting peptides were fingerprinted using the AmaZon ETD ion trap and tandem mass spectrometer prior to shortlisting (Figure 1, A and B, and Figure S1). Five proteins with established roles related to subcellular trafficking, endocytosis, PM targeting and/or endosomal transport were selected (Figure S1 and table S1) and underwent an initial endoribonuclease-prepared siRNA (esiRNA) screen to investigate their impact upon NIS function (Figure 1, C and D). An important finding was that ARF4 depletion repressed iodine-125 (RAI; ^125^I) uptake in MDA-MB-231 NIS+ve cells, whereas VCP ablation resulted in a significant induction of radioiodine uptake (Figure 1D). As a positive control, esiRNA knockdown of PBF, the only protein known to specifically modulate NIS subcellular localisation and function (*36*), significantly increased ^125^I uptake (Figure 1, C and D). In thyroidal TPC-1 NIS+ve cells, treatment with ARF4, VCP or PBF esiRNA also significantly altered radioiodine uptake (Figure S2A).

**Figure 1.**
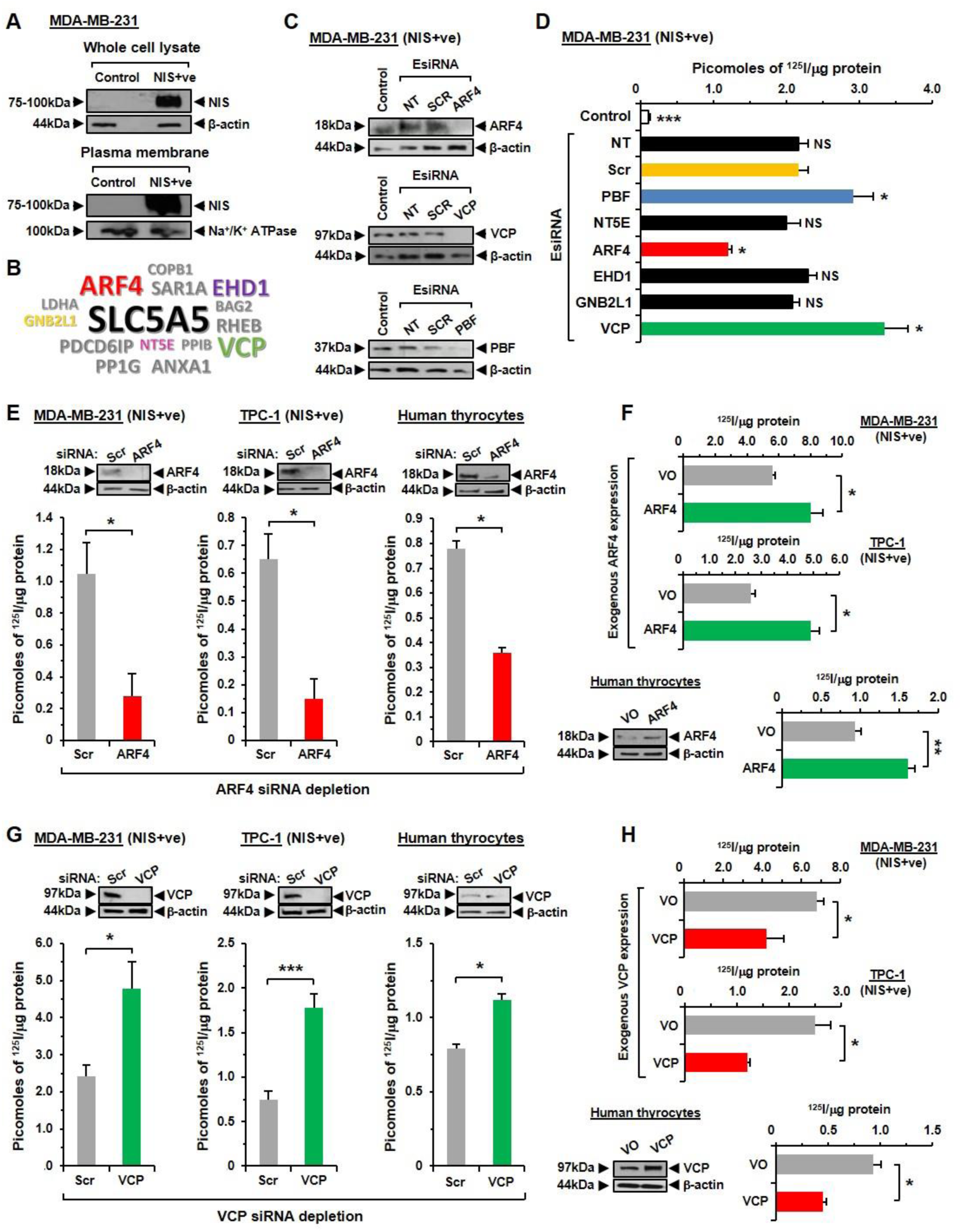
Identification of ARF4 and VCP as regulators of NIS activity. (**A**) Western blot analysis of whole cell lysate and PM fraction in MDA-MB-231 (NIS+ve) cells used in MS/MS to identify NIS interactors. Control – cells transduced with lentiviral (NIS-ve) control particles. (**B**) Top hits for putative NIS interactors identified by MS/MS (peptides≥6). Highlighted proteins were chosen for further study. (**C-D**) Western blot analysis and RAI uptake of MDA-MB-231 (NIS+ve) cells transfected with esiRNA specific for indicated NIS interactors or Scr siRNA. NT – non-transfected cells; Control – same as in (**A**). NS, not significant; *, *P*<0.05; * * *, *P*<0.001, ANOVA with post hoc analysis, n=3. (**E**) RAI uptake in MDA-MB-231 (NIS+ve) cells, TPC-1 (NIS+ve) cells and human primary thyrocytes transfected with ARF4 or Scr siRNA. Representative Western blots of ARF4 protein levels are shown above. (**F**) Same as (**E**) but cells transfected with ARF4 or VO for 48 hours prior to determining RAI uptake. (**G**) Same as (**E**) but cells transfected with VCP or Scr siRNA. Representative Western blots of VCP protein levels are shown above. (**H**) Same as (**G**) but cells transfected with VCP or VO for 48 hours prior to determining RAI uptake. (**E-G**) *, *P*<0.05, * * *, *P*<0.001, student’s t-test. All data presented as mean ± SEM from three independent experiments.

ARF4 and VCP were thus considered novel putative functional partners of NIS and further investigated. Subsequent ARF4 ablation in TPC-1 and MDA-MB-231 NIS+ve cells, as well as in human primary thyroid cells, confirmed a ∼60-80% decrease in ^125^I uptake (Figure 1E) with different siRNA sequences (Table S2). In contrast, transient ARF4 overexpression resulted in significantly increased ^125^I uptake in all three cellular settings (Figure 1F and Figure S2B), suggesting its impact on NIS function is bi-directional. A ∼50-80% increase in ^125^I uptake was further validated in VCP-siRNA-depleted TPC-1 and MDA-MB-231 NIS+ve cells, with human primary thyrocytes demonstrating a ∼35% increase (Figure 1G and table S2). VCP induction by transient transfection reversed these effects, resulting in markedly repressed ^125^I uptake in all three cell models (Figure 1H and Figure S2C).

Thus, we identify two novel modulators of NIS function which alter radioiodine uptake both in thyroid and breast cells lentivirally-transduced with NIS, and in human primary thyrocytes with endogenous NIS expression.

### ARF4 and VCP bind NIS *in vitro* and modulate its expression

Having identified that manipulation of ARF4 and VCP expression altered radioiodine uptake, we sought to challenge our MS/MS data which indicated specific binding between each protein and NIS. Co-immunoprecipitation (Co-IP) confirmed that NIS specifically interacts with ARF4 and VCP in both MDA-MB-231 and TPC-1 NIS+ve cells (Figure 2, A and B). Proximity ligation assays (PLA) further demonstrated specific binding between ARF4 and NIS in three different cell types, which appeared to occur generally throughout the cytoplasm with some binding within the ER/Golgi also suggested (Figure 2C and Figure S3, A and B). Similarly, PLA confirmed the VCP: NIS interaction, which occurred more generally within the cytoplasm of MDA-MB-231, TPC-1 and HeLa cells (Figure 2C and Figure S3, A and B).

**Figure 2.**
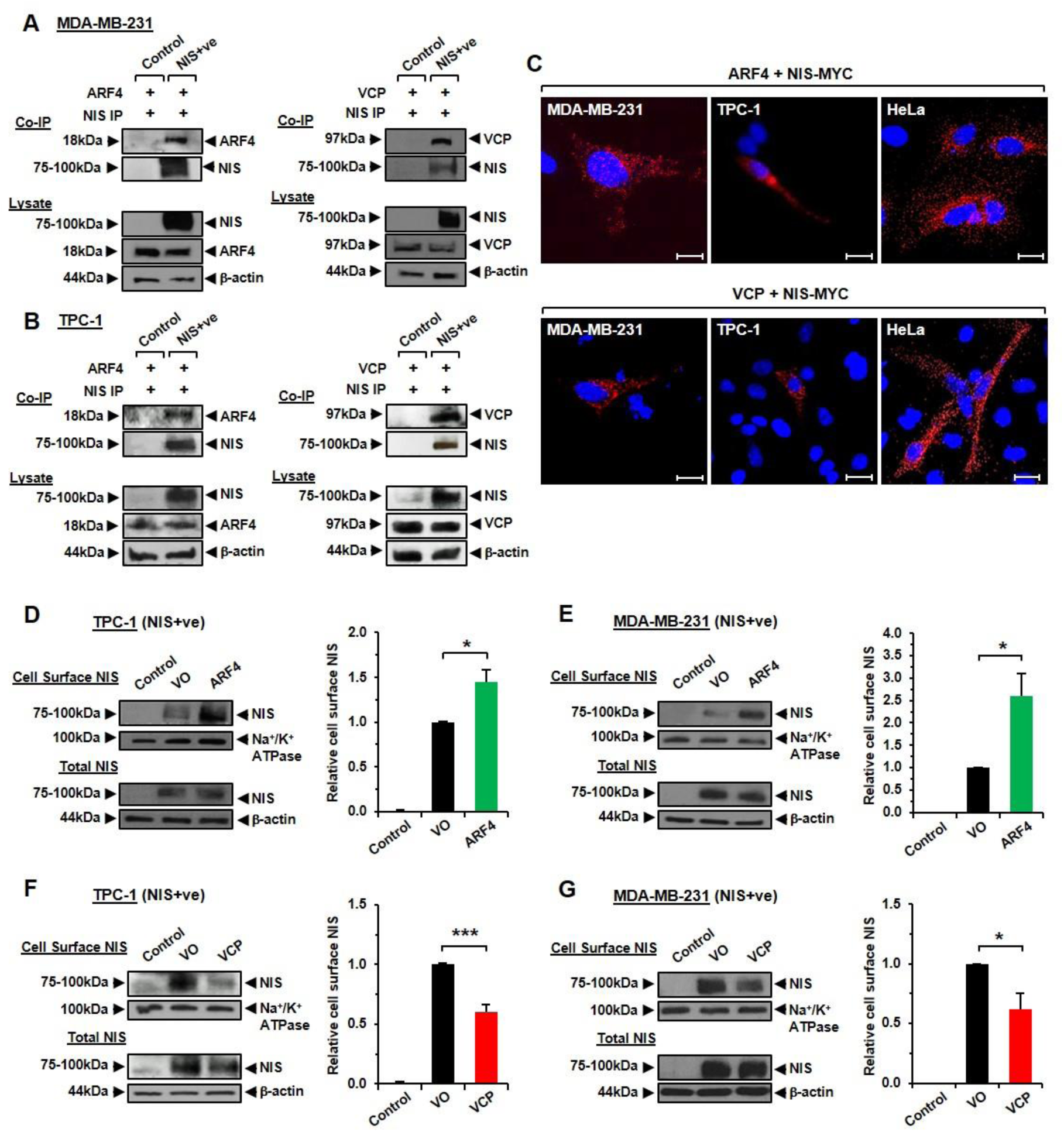
ARF4 and VCP bind NIS *in vitro* and modulate PM NIS. (**A**) Co-IP assays in MDA-MB-231 (NIS+ve) cells showing specific interaction between NIS with ARF4 (left) or VCP (right). Control - cells transduced with lentiviral (NIS-ve) control particles. (**B**) Same as (**A**) but specific interactions shown in TPC-1 (NIS+ve) cells. (**C**) PLA assays showing specific interaction between NIS-MYC and either ARF4 (upper) or VCP (lower) in MDA-MB-231, TPC-1 and HeLa cells. Red fluorescent spots indicate specific interactions. Blue indicates DAPI nuclear staining. Magnification, 100X. Scale bars, 10 μM. (**D-E**) Western blot analysis of CSBA showing that ARF4 overexpression increases NIS protein levels at the PM in TPC-1 (**D**) and MDA-MB-231 (NIS+ve) cells (**E**). Right, mean NIS protein levels at the PM relative to Na^+^/K^+^ ATPase levels. Data presented as mean NIS levels ± SEM from three independent experiments. *, *P*<0.05. (**F**-**G**) Same as (**D**-**E**) but showing that VCP overexpression decreases NIS protein levels at the PM. Right, mean NIS protein levels at the PM relative to Na^+^/K^+^ ATPase. *, *P*<0.05; * * *, *P*<0.001, ANOVA with post hoc analysis. All data presented as mean ± SEM from three independent experiments.

We next performed cell surface biotinylation assays (CSBA) to quantify whether ARF4 and VCP modulate the amount of NIS present at the PM. Exogenous expression of ARF4 in NIS+ve cells demonstrated an approximate doubling of NIS protein in PM preparations compared to VO controls (Figure 2, D and E). In contrast, VCP overexpression resulted in significantly reduced NIS protein PM abundance (Figure 2, F and G) relative to VO-transfected cells. Together, these data suggest that whilst ARF4 potentiates NIS presence at the PM, VCP inhibits its expression at its key site of symporter activity.

### ARF4 modulates NIS trafficking at the PM

Given that ARF4 and VCP overexpression were both associated with altered NIS PM expression, we next investigated their trafficking using HiLo microscopy. We identified that ARF4-dsRED and NIS-GFP trafficked in co-incident vesicles at the PM in HeLa cells (Figure 3, A and B and Figure S4, A and B; Movie S1). By contrast, no such co-trafficking was apparent for VCP-dsRED and NIS-GFP, suggesting the site of functional interaction between VCP and NIS was distant to the PM (Figure S4C and Movie S2). Of particular significance, in contrast to VCP, the presence of ARF4 led to an overall induction in both the mean velocity and distance travelled of NIS-GFP positive vesicles (*P* < 0.001; Figure 3, C and D and Figure S5, A to C).

**Figure 3.**
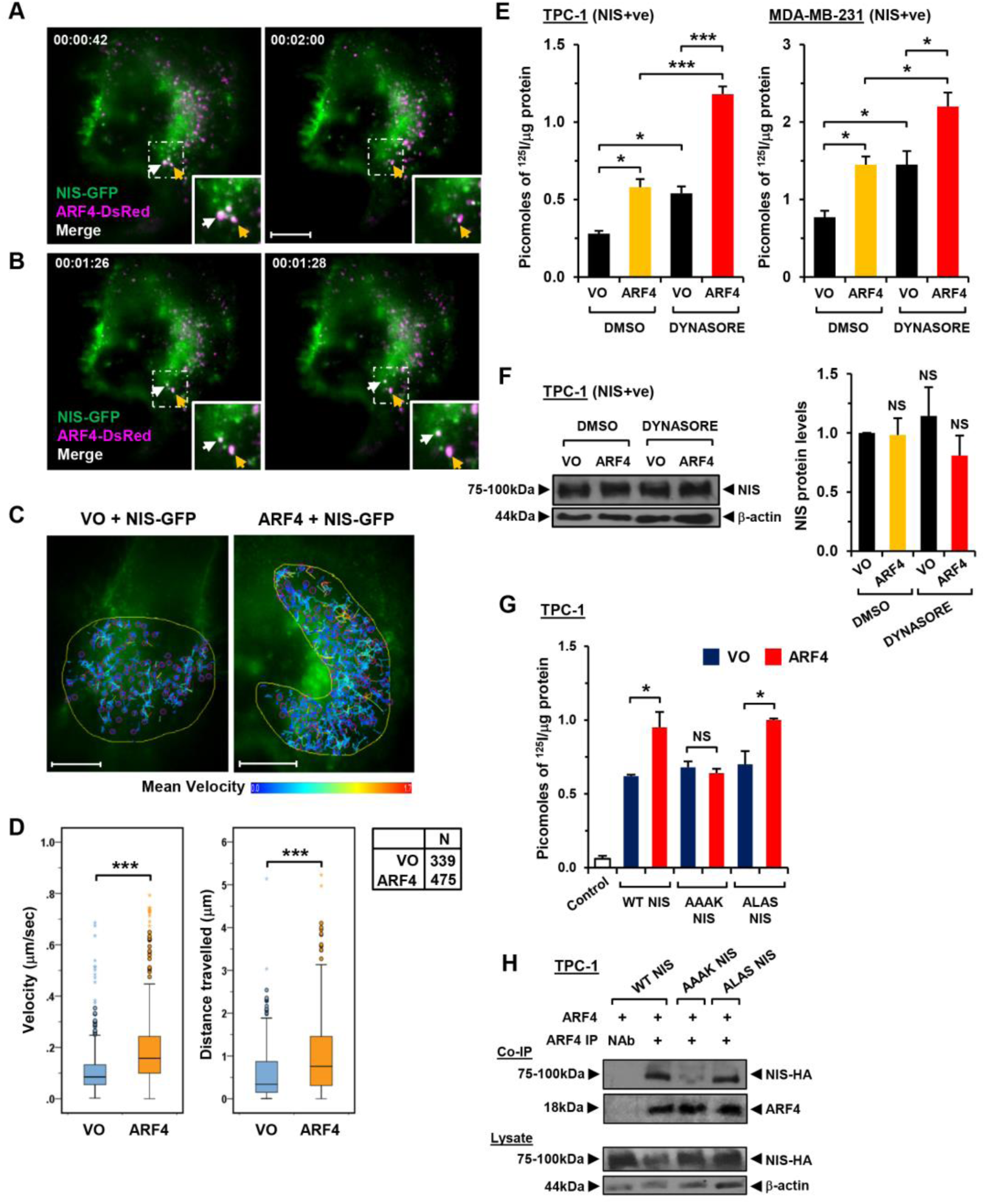
Involvement of ARF4 in trafficking NIS at the PM. (**A-B**) HiLo microscopy images demonstrating trafficking of ARF4-dsRED (magenta), NIS-GFP (cyan) and co-localization (white) to the PM in HeLa cells. Video frame capture times (hr:min:sec) are indicated. PM regions in framed areas are magnified in bottom-right panels which highlight the relatively rapid movement of ARF4 and NIS (white and orange arrowheads; see lower panels). Scale bars, 10 μM. (**C**) Representative images of NIS-GFP movement patterns tracked using ImageJ software. Scale bars, 10 μM. (**D**) Box-whisker plot of velocity (μm/sec) and distance travelled (μm) of NIS-GFP in HeLa cells transfected with ARF4 (*n*=475) or VO (*n*=339). (**E**) RAI uptake in TPC-1 (NIS+ve) and MDA-MB-231 (NIS+ve) cells transfected with ARF4 or VO for 48 hours and treated with Dynasore or DMSO for 1 hour prior to addition of ^125^I. *, *P*<0.05; * * *, *P*<0.001, ANOVA with post hoc analysis. (**F**) Western blot analysis of NIS expression levels in TPC-1 (NIS+ve) cells as described in (**E**). Right, mean NIS protein levels relative to β-actin from three independent experiments. NS, not significant. (**G**) RAI uptake in TPC-1 cells transfected with ARF4 (red) or VO (blue), as well as WT NIS, _574_AAAK_577_ mutant NIS or _475_ALAS_478_ mutant NIS as indicated. NS, not significant; *, *P*<0.05. (**H**) Representative co-IP assay showing that ARF4 binds the _475_ALAS_478_ NIS mutant avidly, whereas the _574_AAAK_577_ NIS mutant abolishes the specific interaction. NAb – no antibody control. All data presented as mean ± SEM from three independent experiments, unless otherwise stated.

In control experiments, we confirmed that NIS is endosomally trafficked in association with clathrin (Figure S5D and Movie S3) and found a significant increase in the overall distance travelled by NIS-GFP in cells overexpressing clathrin (Figure S5A). We then appraised the impact of inhibiting clathrin-mediated endocytosis on NIS function using Dynasore. Treatment of NIS+ve cells overexpressing ARF4 with Dynasore resulted in significantly greater radioiodine uptake (Figure 3E) without any effect on total NIS protein levels (Figure 3F and Figure S5E), suggesting that ARF4 is not primarily enhancing NIS recycling to the PM.

### ARF4 binds NIS via a C-terminal VXPX motif

ARF4 binds to the VXPX motif of Rhodopsin, which represents its four terminal amino acids (*44*). We identified two putative VXPX motifs within the extracellular loop of NIS at positions 475-478 (sequence VLPS) and in the NIS C-terminus at positions 574-577 (sequence VAPK). Abrogation of both motifs (mutations _475_ALAS_478_ and _574_AAAK_577_) resulted in NIS proteins which retained endogenous functionality, but the _574_AAAK_577_ NIS mutant could no longer be augmented in terms of radioiodine uptake by ARF4 overexpression in TPC-1 thyroid cells (Figure 3G). Co-IP assays demonstrated that _475_ALAS_478_ mutant NIS still avidly bound ARF4, whereas _574_AAAK_577_ mutant NIS lost interaction (Figure 3H). Hence, NIS has a VXPX ARF4 recognition sequence in its C-terminus which is required for ARF4 potentiation of function.

Overall, our results indicate that ARF4 and VCP appear to have different modes of action; ARF4 is implicated in the trafficking of NIS to the PM, whereas VCP binds NIS and modulates NIS function elsewhere in the cell. In support of this, significantly altered total NIS protein levels were apparent in NIS+ve cells after siRNA ablation or exogenous expression of VCP but not following modulation of ARF4 expression (Figure S6, A and B).

### Selective VCP inhibitors promote radioiodine uptake

VCP functions chiefly as a chaperone in disassembling protein complexes and facilitating the extraction and/or proteasomal degradation of proteins from the endoplasmic reticulum (ER), but is also implicated in a wider range of other cellular actions (*45*). Several specific VCP inhibitors already exist which target different facets of VCP structure or activity (*46*). Critically, a significant induction of radioiodine uptake (>2.8-fold, Figure 4A) was evident in NIS+ve cells treated with the allosteric VCP inhibitor Eeyarestatin-1 (ES-1), which inhibits ER-cytosol dislocation and subsequent degradation of substrates (*47, 48*). Similarly, the VCP inhibitor NMS-873, which is a potent and specific allosteric inhibitor of VCP capable of activating the unfolded protein response and inducing cancer cell death (*49*), yielded a significant 3-4-fold increase in radioiodine uptake in NIS+ve cells (Figure 4B).

**Figure 4.**
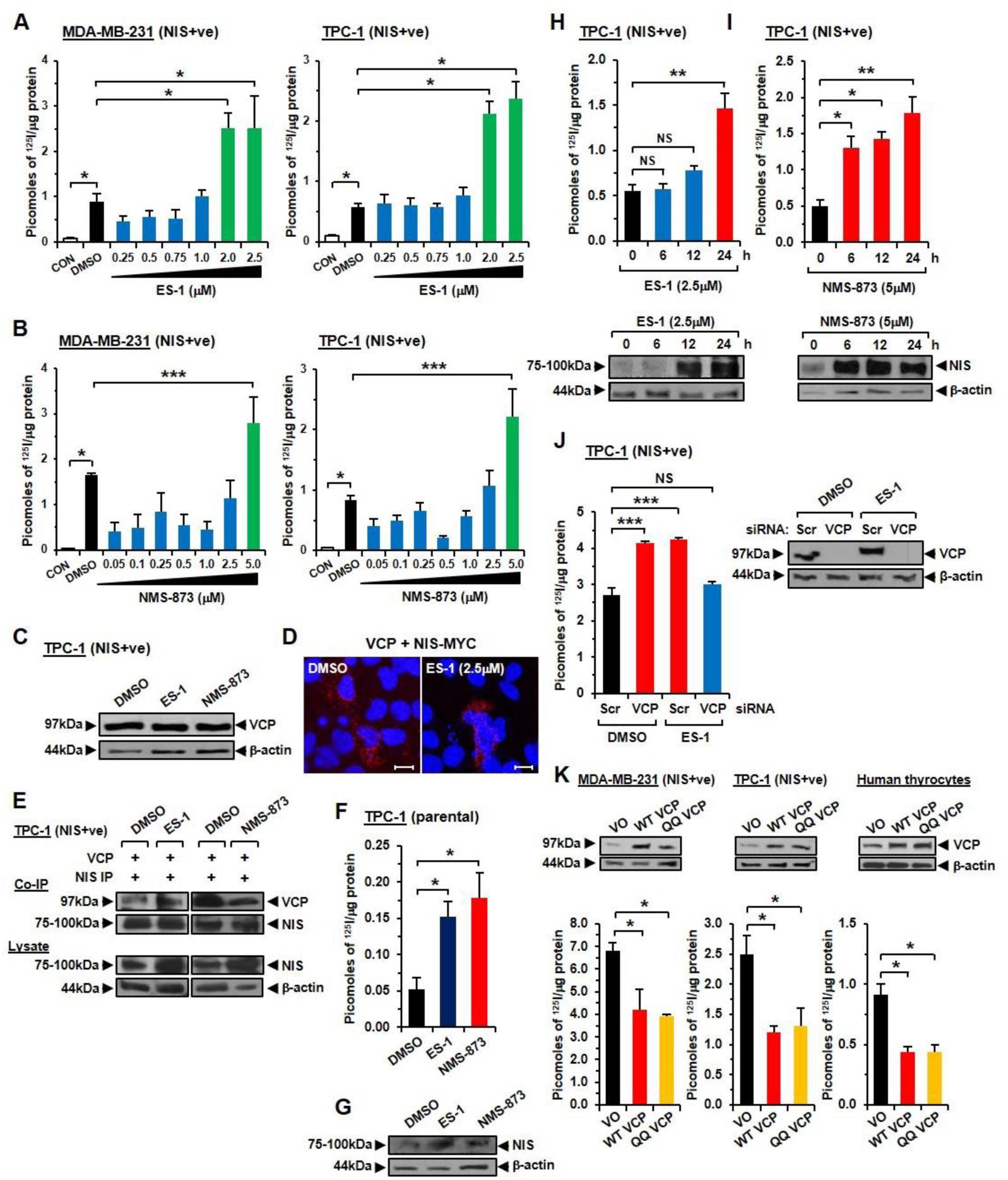
Inhibition of VCP enhances NIS function. (**A**) RAI uptake of MDA-MB-231 (NIS+ve) and TPC-1 (NIS+ve) cells treated with ES-1 at stated doses or DMSO for 24 hours prior to addition of ^125^I. CON - cells transduced with lentiviral (NIS-ve) control particles. (**B**) Same as (**A**) but cells treated with NMS-873. (**C**) Western blot analysis showing that inhibitors ES-1 and NMS-873 do not alter VCP protein expression in TPC-1 (NIS+ve) cells. (**D**) PLA assays showing specific interaction (red fluorescent spots) between NIS-MYC and VCP in TPC-1 cells treated with ES-1 or DMSO. Blue indicates DAPI nuclear stain. Magnification, 100X. Scale bars, 10 μM. (**E**) Co-IP assays showing interaction of VCP with NIS in TPC-1 (NIS+ve) cells treated with ES-1, NMS-873 or DMSO as indicated. (**F-G**) RAI uptake and relative NIS protein levels in parental TPC-1 cells treated with ES-1, NMS-873 or DMSO for 24 hours prior to addition of ^125^I. (**H**) Time-course of RAI uptake and relative NIS protein levels (lower) in TPC-1 (NIS+ve) cells treated with 2.5 μM ES-1 at indicated times (hours) prior to addition of ^125^I. (**I**) Same as (**H**) but cells treated with 5 μM NMS-873. (**J**) RAI uptake in TPC-1 (NIS+ve) cells transfected with VCP or Scr siRNA, and then treated with ES-1 or DMSO as indicated. Western blot analysis (right) confirms VCP ablation in cells transfected with VCP siRNA. (**K**) RAI uptake in MDA-MB-231 (NIS+ve) cells, TPC-1 (NIS+ve) cells and human primary thyrocytes transfected with WT VCP, QQ VCP mutant or VO. Representative Western blots of VCP protein levels are shown above. All data presented as mean ± SEM from three independent experiments. NS, not significant; *, *P*<0.05. * * *, *P*<0.001, ANOVA with post hoc analysis.

Importantly, there were no significant reductions in cell viability at the optimal doses of VCP inhibitors used (Figure S6C), nor any changes in VCP protein expression (Figure 4C). PLA and co-IP assays also revealed that NIS and VCP retained the ability to bind in the presence of VCP inhibitors (Figure 4, D and E). Notably, we also observed a significant induction in radioiodine uptake after 24 hours of ES-1 and NMS-873 inhibitor treatment on native thyroidal cells (i.e. without lentiviral NIS) including TPC-1 (Figure 4, F and G) and Cal62 (Figure S6D). This important finding raises the possibility that VCP inhibitors can be used to induce radioiodine uptake in thyroid cells that have inherently repressed NIS function.

### Dissecting the VCP mechanism of action

We next examined the dynamics of therapeutically targeting VCP function and found that ES-1 was associated with increased detectable NIS protein after 12 hours, accompanied by a concomitant increase in radioiodine uptake by 24 hours (Figure 4H). Similar data were apparent for NMS-873, although radioiodine uptake and NIS expression were detected at 6 hours post-treatment (Figure 4I). To explore the dependency of ES-1 on the presence of VCP to modulate radioiodine uptake, we characterized VCP-ablated TPC-1 NIS+ve cells which demonstrated no induction of radioiodine uptake after ES-1 (Figure 4J) or NMS-873 treatment (Figure S7A). Thus, confirming that inhibitors ES-1 and NMS-873 require VCP in order to exert their effects on NIS function.

VCP’s canonical function lies in facilitating the extraction of misfolded proteins from the ER, a ‘segregase’ process which generally requires ATPase activity. We next used an ATPase-deficient dominant-negative VCP mutant (QQ VCP) (*50*) to investigate whether VCP required ATPase activity to modulate NIS function. The QQ VCP mutant however behaved identically to wild type VCP (WT VCP) in repressing radioiodine uptake when overexpressed in multiple cell models (Figure 4K) and retained the ability to decrease NIS localisation at the PM in CSBA (Figure S7B). Additionally, ATP-competitive VCP inhibitors N2,N4-dibenzylquinazoline-2,4-diamine (DBeQ) (*51, 52*) and sorafenib, which inhibits VCP activity at the PM by blocking its phosphorylation (*53*), failed to increase radioiodine uptake in NIS+ve cells (Figure S7, C and D). Collectively, these results suggest that VCP functions in an ATPase-independent manner to affect NIS function, a facet which has been reported in VCP’s ability to unfold proteins for subsequent proteasomal degradation (*54*).

Recently, three new drugs – astemizole, clotrimazole and ebastine – have been identified as re-purposed small molecules which specifically and allosterically inhibit VCP activity (*55*). Clotrimazole and ebastine are well tolerated *in vivo* and FDA-approved (*55*). As ES-1 and NMS-873 may have limited clinical utility (*56*), we investigated whether astemizole, clotrimazole and ebastine also enhance radioiodine uptake. All three drugs significantly induced radioiodine uptake (Figure 5A) without affecting cell viability (Figure S6C). There was no induction of radioiodine uptake after treatment of VCP-ablated TPC-1 NIS+ve cells, thus confirming these VCP inhibitors require VCP in order to exert their effect on NIS function (Figure 5, B and C). Additionally, VCP inhibitors increased NIS expression, particularly at the PM (Figure 5, D and E), but did not alter VCP protein levels (Figure 5F).

**Figure 5.**
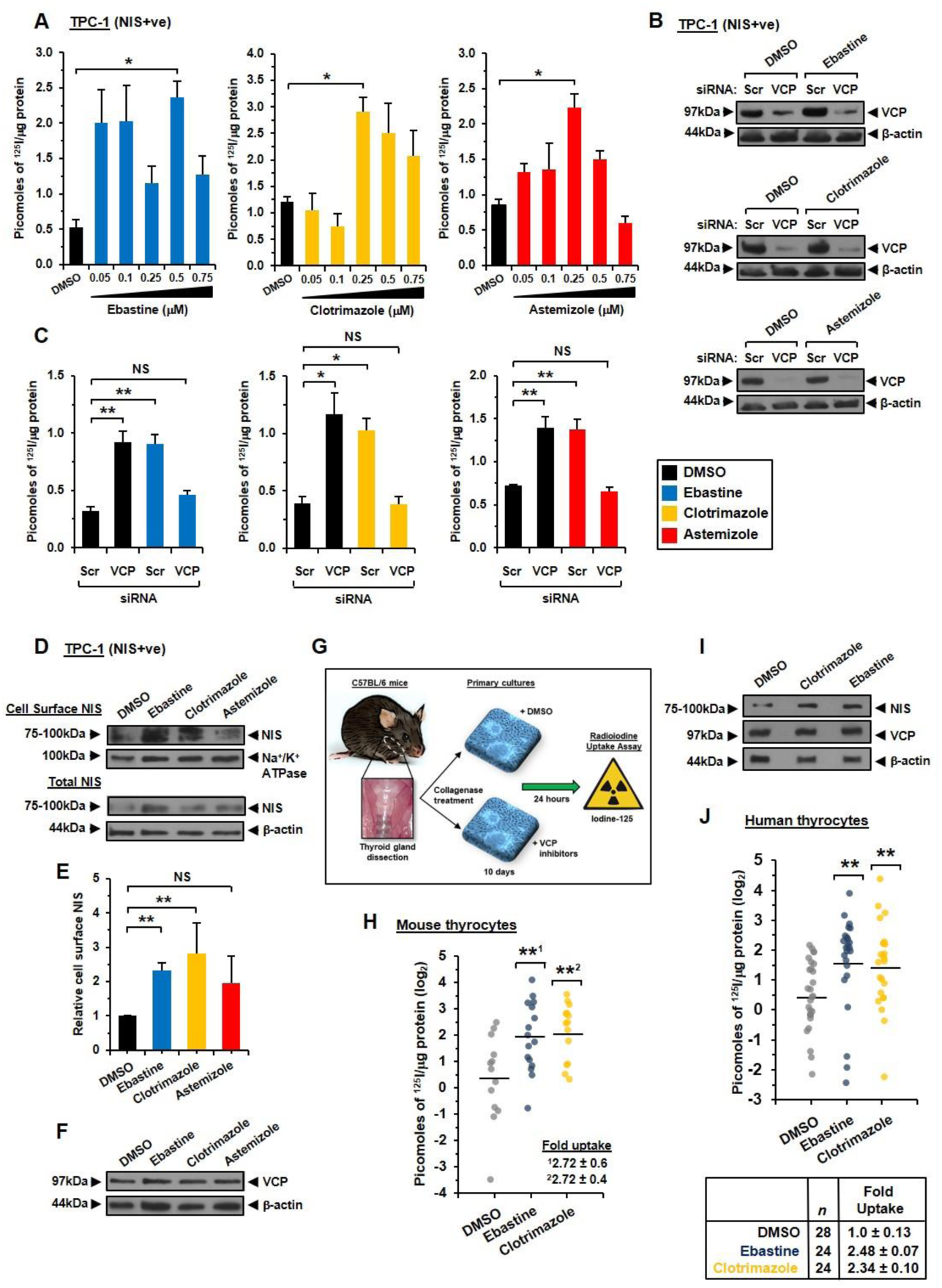
VCP inhibitors increase PM NIS expression and function. (**A**) TPC-1 (NIS+ve) cells treated with stated doses (μM) of VCP inhibitors (Ebastine, Clotrimazole and Astemizole) or DMSO for 24 hours prior to the addition of ^125^I. *, *P*<0.05. * *, *P*<0.01, ANOVA with post hoc analysis. (**B**) Western blot analysis of VCP expression following VCP-siRNA depletion and treatment with 0.5 μM Ebastine (upper), 0.25 μM Clotrimazole (middle), 0.25 μM Astemizole (lower) or DMSO. (**C**) RAI analysis in TPC-1 (NIS+ve) cells transfected and treated as described in (**B**). (**D**) CSBA analysis by Western blot to determine PM NIS expression in the TPC-1 (NIS+ve) cell line 24 hours after treatment with 0.5 μM Ebastine, 0.25 μM Clotrimazole, 0.25 μM Astemizole or DMSO. (**E**) Quantification of Western blots using densitometry to assess PM NIS expression relative to Na+/K+ ATPase. *, *P*<0.05; * *, *P*<0.01. (**F**) Western blot analysis of VCP expression in TPC-1 (NIS+ve) cells treated as described in (**D**). (**G**) Schematic indicating attainment of mouse thyrocytes. Thyroid glands were dissected from C57BL/6 mice, collagenase treated and grown in primary culture then treated with VCP inhibitors for 24 hours prior to the addition of ^125^I. (**I**) RAI analysis of endogenous NIS function in mouse thyrocytes treated for 24 hours with 0.5 μM Ebastine (*n*=8), 0.25 μM Clotrimazole (*n*=8) or DMSO (*n*=9) prior to the addition of ^125^I. (**I**) Western blot analysis in mouse thyrocytes treated as described in (**H**). (**J**) RAI analysis of endogenous NIS function in human thyrocytes treated for 24 hours with Ebastine, Clotrimazole, Astemizole or DMSO prior to the addition of ^125^I. Data presented as maximal response in RAI uptake following treatment with a range of doses of VCP inhibitors due to previous evidence of variability in VCP inhibitor sensitivity (*79*). All data presented as mean ± SEM from three independent experiments, unless otherwise stated.

We next progressed to our model of mouse thyroid function. Primary thyrocytes were isolated from C57BL/6 mice and treated with ebastine or clotrimazole; both drugs significantly enhanced radioiodine uptake and NIS protein expression (Figure 5, G to I). Finally, we tested our drugs in human primary thyroid cultures, which were confirmed TSH-responsive. Both ebastine and clotrimazole significantly increased radioiodine uptake (Figure 5J).

### VCP and ARF4 expression correlates with poorer survival and response to RAI

Having identified VCP and ARF4 as novel regulators of NIS function, we appraised their expression profiles and clinical relevance in thyroid cancer (THCA) via The Cancer Genome Atlas database (TCGA). Of significance, ARF4 was under-expressed in PTC compared to normal tissue (Figure 6A), while VCP was significantly induced in PTC (Figure 6A). Genetic drivers in PTC have distinct signalling consequences and have been categorized into BRAF-like and RAS-like PTCs according to distinct gene signatures (*57*). Interestingly, VCP mRNA expression did not differ between BRAF-like and RAS-like PTC, while ARF4 mRNA expression was lower in BRAF-like than RAS-like tumours across the THCA series (*n*=391; Figure 6B).

**Figure 6.**
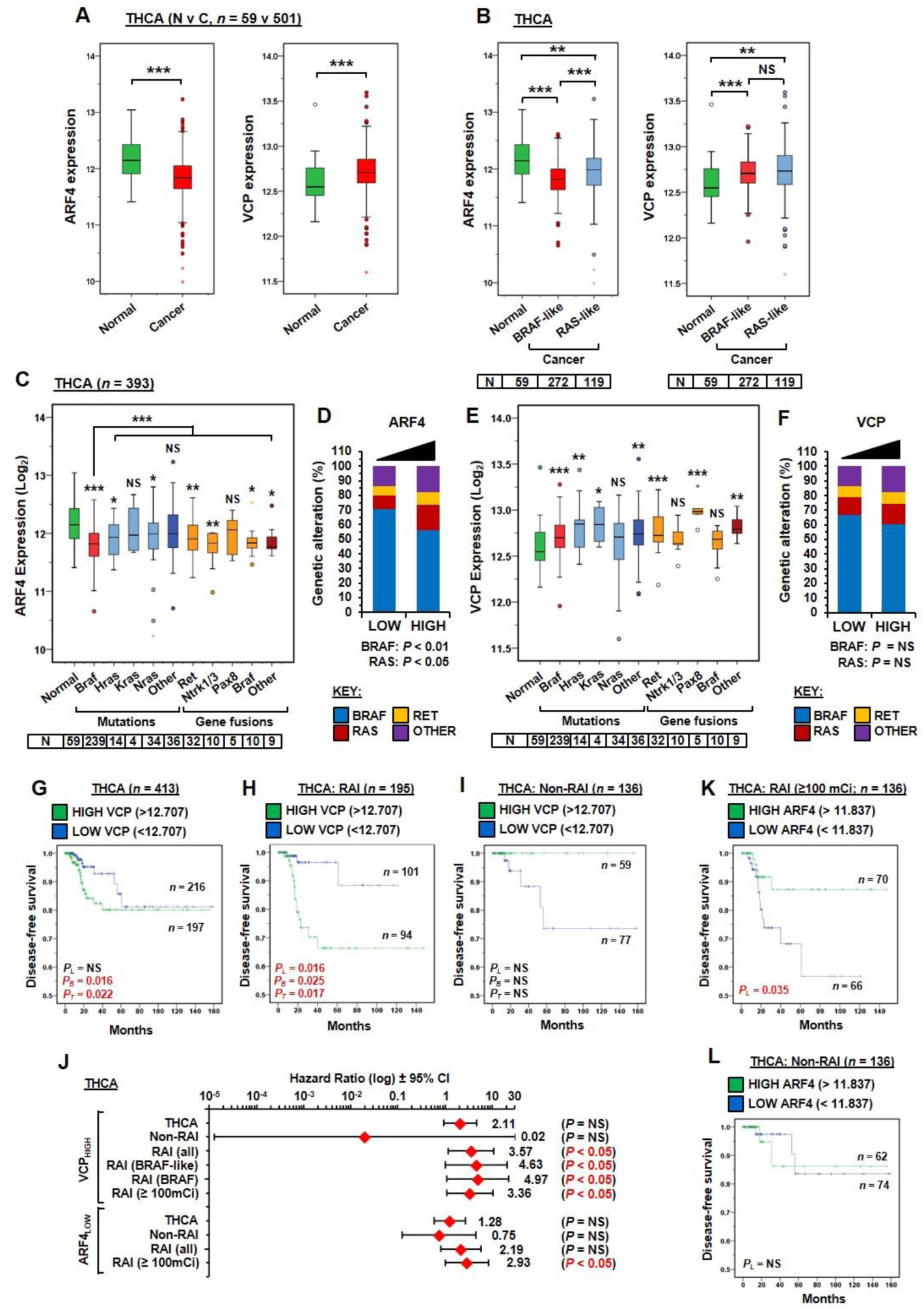
VCP and ARF4 expression associates with poorer survival and response to RAI. (**A-B**) Box whisker plots showing ARF4 (left) and VCP (right) expression in normal (*n*=59) versus cancer (*n*=501) in THCA TCGA data (**A**), and in PTC with either BRAF-like or RAS-like genetic signatures relative to normal (**B**). NS, not significant; * *, *P*<0.01; * * *, *P*<0.001. (**C**) Box-whisper plots of ARF4 expression in PTC with the indicated genetic alteration and number of samples relative to normal. NS, not significant; *, *P*<0.05; * *, *P*<0.01; * * *, *P*<0.001. (**D**) Frequency (%) of indicated genetic alterations in PTC with low (Q1Q2) versus high ARF4 (Q3Q4) expression. (**E**-**F**) Same as (**C**-**D**) but showing THCA TCGA data for VCP expression. (**G-I**) DFS for THCA with high (Q3Q4) versus low (Q1Q2) VCP expression for the entire PTC cohort (**G**), RAI-treated patients (**H**) and non-RAI treated patients (**I**). (**J**) Hazard ratios ±95% CI for patients stratified on median VCP and ARF4 tumoural expression in THCA with the indicated treatment and genetic signature or alteration. VCP_HIGH_ or ARF4_LOW_ expression was associated with significantly increased risk of recurrence/relapse as indicated. (**K-L**) DFS for THCA with high (Q3Q4) versus low (Q1Q2) ARF4 expression for RAI-treated PTC patients (**K**) and non-RAI treated PTC patients (**L**).

In agreement with these findings, there was greater reduction of ARF4 expression in BRAF-mutant PTC versus the non-BRAF mutant tumours (Figure 6C). The frequency of *BRAF* alterations was also higher in PTC (70.6%; *n*=139/197) with low ARF4 expression compared to PTC with high ARF4 (56.1%; *n*=110/196; Figure 6D). By comparison, VCP expression was most significantly elevated in PTC with *BRAF* mutant, *RET* fusion and *PAX8* fusion genes (Figure 6E). In contrast to ARF4, there was a similar frequency of *BRAF* alterations in PTC with either high or low VCP expression (Figure 6F).

Survival profiles in THCA are available for PTC patients treated with a range of radioiodine (RAI) doses, as well as those who have not received any radioiodine (i.e. non-RAI). We therefore evaluated whether VCP and ARF4 expression were associated with patient outcome and treatment. Overall, log-rank analysis using the entire cohort (*n*=413) did not detect any difference in survival between PTC patients with high tumoural VCP compared to those with low VCP (*P*_*L*_=NS), although a significant difference was evident at early (*P*_*B*_=0.016) and intermediate time-points (*P*_*T*_=0.022; Figure 6G). Similarly to THCA, higher VCP expression was present in breast tumours versus normal tissue in the BRCA cohort (Figure S8A), which again failed to correlate with a significant reduction in survival (*P*_*L*_=NS; Figure S8B).

A key finding however was that the subgroup of RAI-treated PTC patients with high tumoural VCP expression (*n*=195) had significantly reduced disease-free survival (DFS) than those with low VCP (*P*_*L*_=0.016; Figure 6H and Figure S8C). By comparison, there was no significant difference in DFS of patients that did not receive RAI treatment when stratified on median tumoural VCP (Figure 6I). Cox regression analysis further highlighted that higher tumoural VCP in RAI-treated PTC patients was associated with an increased risk of recurrence (HR 3.57, 95% CI: 1.18 – 10.86; Figure 6J). Interestingly, the risk of recurrence was even greater for RAI-treated patient subgroups associated with BRAF-like (HR 4.63; 95% CI: 3.61 – 16.31) or BRAF alterations (HR 4.97; 95% CI: 3.87 – 17.47; Figure 6J and Figure S8, D to G). RAI-treated patients with lower tumoural ARF4 expression receiving a dose ≥100mCi also had reduced survival (Figure 6, K and L, and Figure S8, H and I) and a higher risk of recurrence (HR 2.93, 95% CI 1.03 – 8.33; Figure 6J). In contrast, there was no increase in the risk of recurrence of non-RAI treated PTC patients with higher tumoural VCP or lower ARF4 expression (Figure 6J).

Collectively, we show that ARF4 and VCP are significantly dysregulated in PTC, with expression profiles which fit the repression of radioiodine uptake generally apparent in thyroid cancers. We propose that dysregulated ARF4 and VCP results in reduced trafficking of NIS to the PM via repressed ARF4 function and increased VCP activity in targeting NIS for degradation. We further identify that VCP and ARF4 are associated with poorer survival characteristics in RAI-treated patients and represent promising new drug targets in patients who are radioiodine-refractory. A model of the proposed functional interaction of NIS with VCP and ARF4 is outlined in Figure 7.

**Figure 7.**
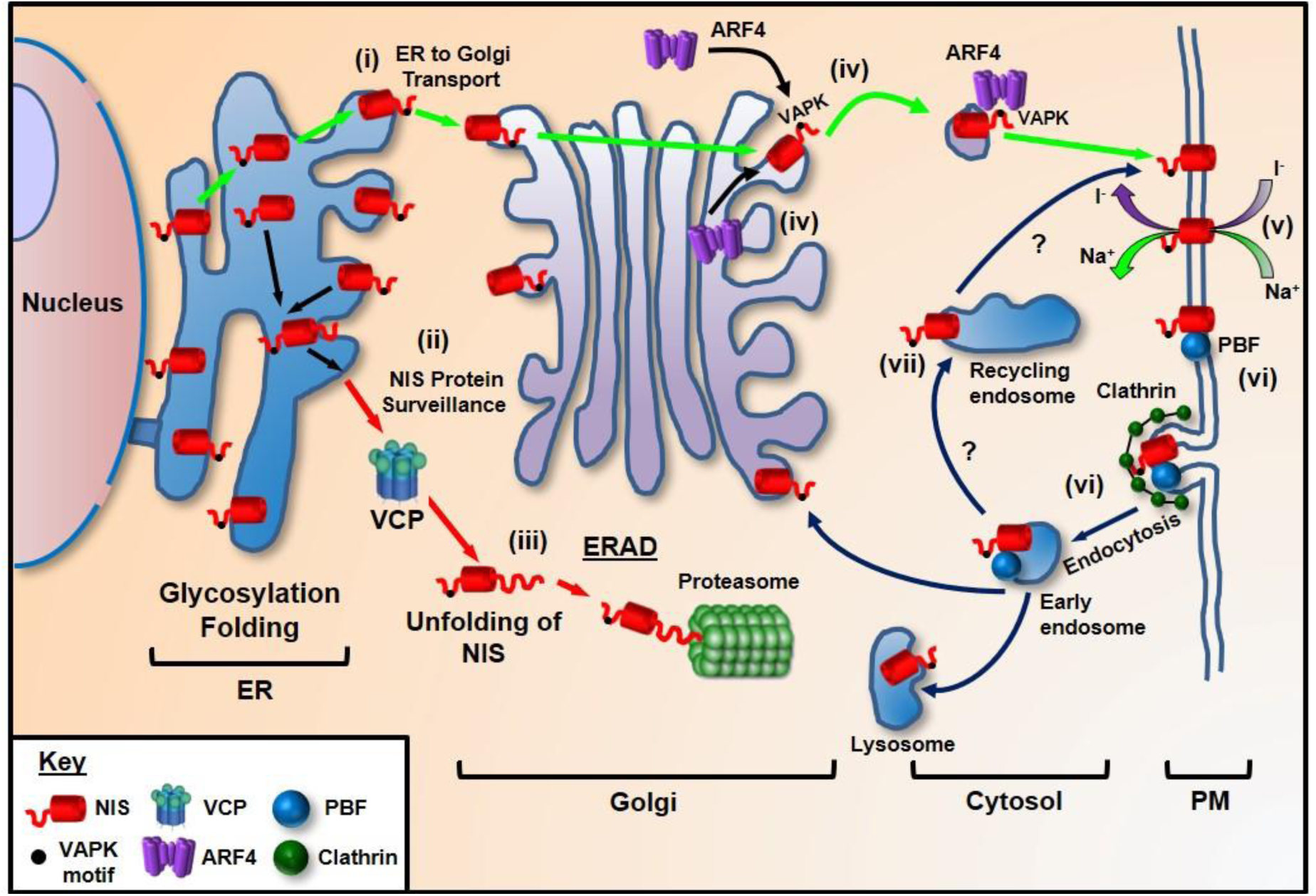
Putative model of NIS trafficking. NIS maintains a delicate balance between protein synthesis, folding, assembly, trafficking and degradation. (**i**) We predict that NIS is glycosylated in the endoplasmic reticulum (ER) and upon correct folding transported to the Golgi. (**ii**) Protein surveillance pathways exist that target NIS for ERAD. As VCP does not require ATPase activity to inhibit NIS function, it is likely that VCP acts to unfold NIS (**iii**) prior to proteasomal degradation, which, due to the constraint of pore size, would require NIS to be unstructured. (**iv**) ARF4 recognises the VAPK residues of the NIS C-terminus and promotes vesicular trafficking to the PM, where NIS is active (**v**). (**vi**) PBF has a YARF endocytosis motif and acts to bind and internalise NIS away from the PM in a clathrin-dependent process. (**vii**) Although inhibition of recycling by Dynasore suggests that ARF4 shuttles NIS to the PM, other proteins must promote recycling of NIS to the PM as with the majority of PM transporters.

## DISCUSSION

Extensive studies have sought to enhance NIS expression and function in patients with radioiodine-refractory thyroid cancer (RAIR-TC), and hence re-sensitise tumours to radioiodine therapy. This is essential because patients with RAIR-TC, particularly those with metastatic disease, have a life expectancy of 3–5 years and represent a group for whom there is a clear unmet medical need (*1, 2, 58*). Most investigations so far have focussed on ‘re-differentiation agents’, which stimulate the expression of thyroid-specific genes, including NIS. Constitutive activation of the MAPK pathway in thyroid cancer results in dysregulated NIS expression and function, decreased radioiodine uptake and a poor patient prognosis (*59*). A substantial number of studies have therefore focussed on selective inhibitors of the MAPK pathway to restore radioiodine avidity (*5, 60*).

The lesional radiation dose attained after radioiodine therapy depends on the uptake of ^131^I by thyroid cells via NIS, and its retention time in thyroid tissue. Retention requires iodide oxidation by thyroid peroxidase, contingent upon H_2_O_2_ production by dual oxidases Duox1 and Duox2 (*61*), and subsequent incorporation into thyroglobulin. Importantly, new drug strategies, such as combining BRAF or MEK inhibitors with pan-PI3K inhibitors, are showing pre-clinical promise by rescuing thyroid-specific gene expression and enhancing iodide retention (*25*). However, we propose that augmenting NIS trafficking to the PM is fundamental to boosting the efficacy of radioiodine treatment, given that we are now close to having the necessary tools to restore NIS expression and iodide retention.

Here, we identified five structurally diverse allosteric inhibitors of VCP which all enhanced radioiodine uptake *in vitro* by a minimum of two-fold. Specific ARF4 agonists do not currently exist, and hence we pursued VCP inhibition as our central therapeutic strategy. CB-5083, the first-in-class inhibitor of VCP, was shown to be effective in inhibiting tumour growth in pre-clinical models (*62*). Clinical trials of CB-5083 for solid and haematological malignancies however were recently prematurely terminated due to off-target effects on PDE6 resulting in ocular dysfunction (*63*). To circumvent potential issues of toxicity we evaluated alternative drugs to enhance radioiodine uptake, of which two of them - ebastine and clotrimazole - are well tolerated *in vivo* and FDA-approved (*55*). Given that radioiodine uptake was enhanced within 24 hours of treatment, our findings necessitate the need for clinical trials to address whether patients receiving already well-tolerated doses of ebastine or clotrimazole at the time of radioiodine treatment uptake more ^131^I than those given placebos. One important finding in support of this is that native, relatively de-differentiated thyroid cells which have negligible endogenous radioiodine uptake in our hands showed measurable NIS function following VCP inhibition.

Our MS/MS, co-IP and PLA assays all identified and confirmed ARF4 and VCP as new NIS interactors. The endocytosis of NIS is known to be modulated by the protein PBF (*36*). Previously PBF which, unlike NIS, has a functional endocytosis motif (*64*), was the only protein shown to bind NIS and alter its subcellular localisation (*36*). We now identify the novel NIS interactor ARF4 binds NIS via a C-terminal _574_VXPX_577_ motif to traffic NIS towards the PM and enhance NIS function.

We sought to apply new imaging technologies to address the cellular mechanisms of NIS trafficking. One surprising facet was the rapid trafficking of vesicles expressing both ARF4 and NIS, which hint at a more dynamic process of NIS trafficking than previously suggested. ARF4 has a range of subcellular roles, from recruiting adaptor proteins for packaging proteins into vesicles destined for the PM, to recycling proteins from the PM through endosomes (*65*). Our data suggest ARF4 is involved in the vesicular transport of NIS to the PM, given that inhibition of endocytosis had no impact on the ARF4-mediated increase in radioiodine uptake.

An important clinical observation was that high tumoural VCP expression, which acts predominately in the ER, and hence early in the progression of NIS to the PM, was strongly correlated with DFS, both overall, but particularly in patients who received radioiodine treatment. Similarly, PTC patients with low tumoural ARF4 expression who received higher doses of radioiodine (≥100 mCi) also had a significantly worse DFS. Our hypothesis is that high tumoural VCP expression in patients with PTC results in increased NIS degradation, hence permitting less NIS to be trafficked to the PM by ARF4, which we show is repressed in PTC. Thus, the function of NIS, which is already at very low expression levels or predominantly localized to intracellular compartments in PTC is further attenuated, resulting in worse DFS.

Several gaps in our knowledge remain. As a protein expressed mainly in basolateral membranes, cell polarisation is important to NIS function. There are no human polarised cell models which are physiologically relevant to thyroid cell biology, however. Despite this, our studies in non-polarised cell systems demonstrate a universal mechanism of NIS trafficking in transformed thyroid and breast cells, as well as primary thyrocytes. Further, since our de-differentiated TPC-1 thyroid cells, as well as our MDA-MB-231 breast cells, are unlikely to express physiologically relevant levels of the TSH receptor, we did not investigate the potential effects of TSH – the canonical regulator of NIS – on NIS trafficking (*66*). Our hypothesis is that TSH is more important to the expression of NIS than its direct trafficking. This reflects the surprising timescales of trafficking we observed in our HiLo microscopy; whereas TSH is known to influence NIS function over the course of hours and days (*66*), vesicular movement of NIS close to the PM was unexpectedly dynamic. We thus propose that NIS function at the PM is a much more rapid process than currently envisaged.

Together, we now identify two proteins, VCP and ARF4, with critical roles in NIS function which correlate markedly with clinical outcome in PTC. In particular, VCP is specifically druggable with existing inhibitors resulting in clearly induced radioiodine uptake. Strategies that manipulate the function/expression of VCP or ARF4 will thus offer a promising new therapeutic strategy for RAIR-TC.

## MATERIALS AND METHODS

### Mass spectrometry

NIS-interactors were isolated by co-IP (anti-NIS antibody) and separated by SDS-PAGE. Proteins were reduced, alkylated and trypsinised. Resulting peptides were separated on an acetonitrile gradient on the UltiMate 3000 HPLC (ThermoFisher Scientific). Eluted peptides passed through the AmaZon ETD ion trap and tandem mass spectrometer (Bruker). Mass spectra were processed using the Bruker DataAnalysis software and analyzed using the Mascot search engine (Matrix Science). Datasets were filtered based on peptide number, DAVID functional classification (*67, 68*) and literature review (*69-73*).

### Western blotting, cell-surface biotinylation and co-IP assays

Western blotting and cell surface biotinylation assays (CSBA) were performed as described previously (*36, 74, 75*). Blots were probed with specific antibodies (Table S3) and NIS expression quantified by densitometry in ImageJ relative to β-actin or Na^+^/K^+^ ATPase, as indicated. Co-IPs were performed as described (*36*) except cell lysates underwent Dounce homogenisation to facilitate lysis.

### HiLo microscopy and proximity ligation assay

Live cell HiLo microscopy was performed on an Olympus IX81 inverted fluorescence microscope. Images were acquired every 2 seconds for up to 5 minutes then integrated at 5 frames/second using the Olympus xCellence build 3554 software (see Movies S1 to 3). Mean velocity and distance travelled of NIS-GFP was quantified using the TrackMate plugin (ImageJ). The Duolink® *in situ* proximity ligation assay (PLA) was performed according to the manufacturer’s instructions (Sigma-Aldrich).

### Radioiodine uptake and inhibitors

Radioiodine (RAI; ^125^I) uptake was performed as described previously (*64*) following VCP inhibitor treatment (24 hours), DNA plasmid transfection (48 hours) and siRNA transfection (72 hours) as indicated. In some experiments, cells were transfected with siRNA for 48 hours prior to VCP inhibitor treatment for the remaining 24 hours.

### TCGA data analyses

Normalized gene expression data and clinical information were downloaded from TCGA via cBioPortal (*76, 77*) and FireBrowse (*78*). In total, RNA-seq data for 501 THCA and 1093 BRCA TCGA samples were analyzed.

### Statistical analyses

Data were analyzed using GraphPad Prism and Microsoft Excel. For comparison between two groups, data were subjected to the Student’s t-test and Mann-Whitney U-test, and for multiple comparisons one-way ANOVA with Tukey’s test for multiple comparisons and Kruskal– Wallis tests were used. Primary thyrocyte data underwent log-transformation prior to analysis. For DFS, the median value of VCP and ARF4 expression (log_2_) in TCGA was used to stratify patients with high or low expression. Subsequent analyses were completed using Kaplan-Meier curves and cohort comparisons were performed by the Log rank (P_L_) Breslow (P_B_) and Tarone-Ware (P_T_) tests. All grouped data are presented as mean ± SEM for three independent replicates unless otherwise stated, *P* < 0.05 was considered significant.

## DATA AVAILABILITY

The source data for Figures 1-6, and Supplementary Figures are provided as a Source Data file. Other data are available upon request.

## AUTHOR CONTRIBUTIONS

C.J.M., V.E.S. and A.F. conceptualized/planned the study. A.F., M.L.R., C.E.M.T., D.P.L., V.L.P., K.Br., H.R.N., M.A. and R.J.T. acquired the data. A.F., M.L.R. and M.J.C. analyzed/interpreted data. A.F., M.L.R., G.G.L., K.Bo., V.E.S. and C.J.M. prepared/reviewed the manuscript. K.Bo., A.S.T., V.E.S. and C.J.M. provided supervision.

### ACKNOWLEDGEMENTS

This work was supported by MRC (GBT1529), Wellcome Trust (RCHX21285), Get A-Head and The University of Birmingham. Results are in-part based upon data generated by TCGA: http://cancergenome.nih.gov/. We thank A. Di Maio (University of Birmingham) and D. Calebiro (University of Birmingham) for expertise in HiLo; D. Nasteska (University of Birmingham) and D. Hodson (University of Birmingham) for the C57BL/6 mice. We acknowledge the contribution to this study made by the Human Biomaterials Resource Centre (University of Birmingham).

## COMPETING INTERESTS

The authors declare no competing interests.

