## Supplemental Material for "Pharmacologically targeting a novel pathway of sodium iodide symporter trafficking to enhance radioiodine uptake"

**This file includes:**

**Materials and Methods**

Human thyroid tissue and cell culture

Mouse primary thyrocytes

Nucleic acids and transfection

**Supplementary figures**

Figure S1. Identification and selection of putative NIS interactors.

Figure S2. siRNA ablation of ARF4 and VCP alters $^{125}I$ uptake in thyroid cells.

Figure S3. Novel functional partners ARF4 and VCP specifically interact with NIS.

Figure S4. NIS traffics with ARF4, but not VCP, at the PM.

Figure S5. ARF4 increases the velocity and distance travelled of NIS-GFP positive vesicles.

Figure S6. ARF4 and VCP differentially affect NIS expression, but VCP inhibitors do not alter cell viability.

Figure S7. VCP inhibitors require VCP expression to exert their effect on NIS function.

Figure S8. VCP expression correlates with worse DFS in PTC with BRAF alterations.

Figure S9. Short tandem repeat (STR) profiling of cell lines used in study.

**Supplementary tables**

Table S1. Top hits for putative NIS interactors identified by mass spectrometry.

Table S2. siRNAs used in study.

Table S3. Primary antibodies used in study.

Table S4. Nucleic acids used in study.

**Supplementary movies**

Movie S1: Significant co-localisation and trafficking of ARF4-dsRED and NIS-GFP proteins in co-incident vesicles at the plasma membrane in HeLa cells

Movie S2: Lack of co-trafficking for VCP-dsRED and NIS-GFP suggests that the site of functional interaction between VCP and NIS is distant to the plasma membrane in HeLa cells

Movie S3: The sodium iodide symporter NIS is endosomally trafficked in association with clathrin

### SUPPLEMETNARY MATERIALS

#### Materials and Methods

##### Human thyroid tissue and cell culture

Human thyroid tissue was obtained with local ethics committee approval from the Human Biomaterials Resource Centre (University of Birmingham) and informed patient consent. Primary thyrocytes were isolated and cultured as described (*80*). MDA-MB-231 breast and TPC-1 thyroid cancer cell lines were maintained in RPMI-1640 (Life Technologies), while HeLa cervical cancer cells were maintained in DMEM (Sigma-Aldrich). Media was supplemented with 10% fetal bovine serum (FBS), penicillin ($10^5$ U/l), and streptomycin (100 mg/l) and cell lines were maintained at 37°C and 5% $CO_2$ in a humidified environment. All cell lines were obtained from the ECACC except TPC-1, which were kindly provided by Dr Rebecca Schweppe (University of Colorado). Cells were cultured at low passage, authenticated by short tandem repeat analysis (NorthGene; Figure S9) and tested for mycoplasma contamination (EZ-PCR kit; Geneflow).

Stable NIS-expressing MDA-MB-231 and TPC-1 cell lines were generated by lentiviral transduction, as per manufacturer's instructions. In brief, ready-to-transduce lentiviral particles containing a precision lentiORF construct (pLOC) housing cDNA coding for *RFP* (OHS5833) or the full-length human *NIS* cDNA without a stop codon within the open reading frame (ORF) (OHS5900-224632369) were purchased from Dharmacon. For lentiviral transduction the manufacturer's protocol was followed. In brief, one day after plating, MDA-MB-231 and TPC-1 cells were infected with the lentiviral vector containing *NIS* diluted in antibiotic-free and serum-free RPMI containing 8 µg/ml or 14 µg/ml polybrene respectively. Cells transduced with the *RFP*-containing lentiviral vector served as the control. After 24 hours, medium was replaced with RPMI containing 10% fetal bovine serum (FBS). After a further 48 hours, cells were maintained in RPMI containing 10% fetal bovine serum (FBS), penicillin ($10^5$ U/l), and streptomycin (100 mg/l) as well as 15 or 5 µg/ml *Blasticidin S* for the MDA-MB-231 and TPC-1 cell lines respectively. 24 hours later, infection efficiency was assessed by fluorescence microscopy. Upon cell expansion and selection of single cell colonies, NIS expression was assessed by quantitative PCR (qPCR) and Western blotting, with NIS function confirmed via the radioiodine uptake assay.

**Mouse primary thyrocytes**

C57BL/6 mice were bred at the Biomedical Services Unit (University of Birmingham) and experiments were performed in accordance with UK Home Office regulations. At 6-10 weeks of age, thyroid glands were removed and primary thyrocytes cultured as described previously (*81*).

**Nucleic acids and transfection**

Plasmids containing human NIS cDNA with a HA- or MYC-tag have been described (*64*) and GFP-tagged NIS was kindly provided by Dr Takahiko Kogai (Dokkyo Medical University). NIS mutants $_{475}$ALAS$_{478}$ and $_{574}$AAAK$_{577}$ were generated using the QuikChange Site-directed Mutagenesis Kit (Agilent Technologies). Untagged-ARF4 plasmid was purchased from Origene (#SC119092) while wild type (WT) and mutant (QQ) VCP (rat) plasmid were kindly provided by Dr Yihong Ye (National Institutes of Health). Human VCP cDNA was generated by site-directed mutagenesis. Further details on nucleic acids are provided (Table S4). Plasmid DNA and siRNA transfections were performed with TransIT-LT1 (Mirus Bio) and Lipofectamine RNAiMAX (ThermoFisher Scientific) following standard protocols.

#### Supplementary figures

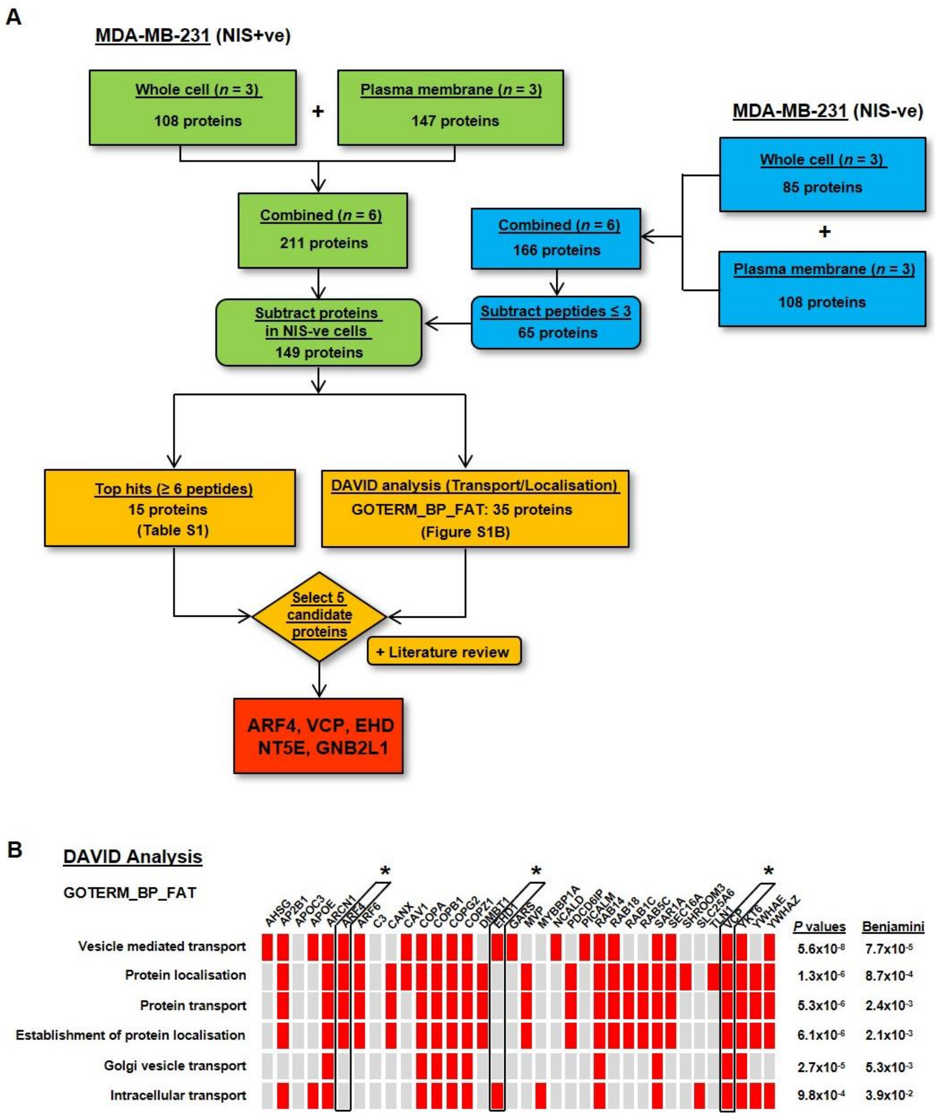


**Figure S1. Identification and selection of putative NIS interactors.** (**A**) MS/MS was performed following co-IP of NIS and its interactors using an anti-NIS antibody in whole cell (*n* = 3) and PM fractions (*n* = 3) extracted from MDA-MB-231 (NIS+ve) and control (NIS-ve) cells. Known atmospheric, experimental and non-specific contaminants were filtered from the dataset for MDA-MB-231 (NIS+ve) cells prior to subtracting 65 proteins with >3 peptides identified in control (NIS-ve) cells. The remaining 149 proteins (see Source Data File) were filtered based on peptide number, DAVID functional classification (*67*, *68*) and literature review (*69*-*73*). Proteins with ≥ 6 peptides were considered top hits (Table S1). Based on this criteria, five candidate genes (i.e. ARF4, VCP, EHD1, NT5E and GNB2L1) were selected for further study. NT5E was shortlisted as it has been previously described as having a role in hypo- and hyper-thyroidism whereby T3 and T4 modulate its expression and function (*69*, *72*), as well as being overexpressed in metastatic disease (*73*). GNB2L1 was shortlisted as it is overexpressed in PTC and has been associated with the BRAF V600E mutation (*71*). GNB2L1 is also involved in membrane trafficking and the facilitation of protein-protein interactions through its role as a scaffold protein (*70*, *71*). (**B**) Functional annotation of the filtered list of 149 proteins (Figure S1A) using the GOTERM_BP_FAT category in DAVID (version 6.7; https://david.ncifcrf.gov/) indicated that 35 proteins had significantly enriched gene ontology (GO) terms associated with either transport or localization (marked in red). Of these, three proteins (i.e. ARF4, EHD1 and VCP) were chosen for further study and are marked with an asterisk. *P* values are from modified Fisher Exact tests. Benjamini values are *P* values corrected for multiple comparisons.

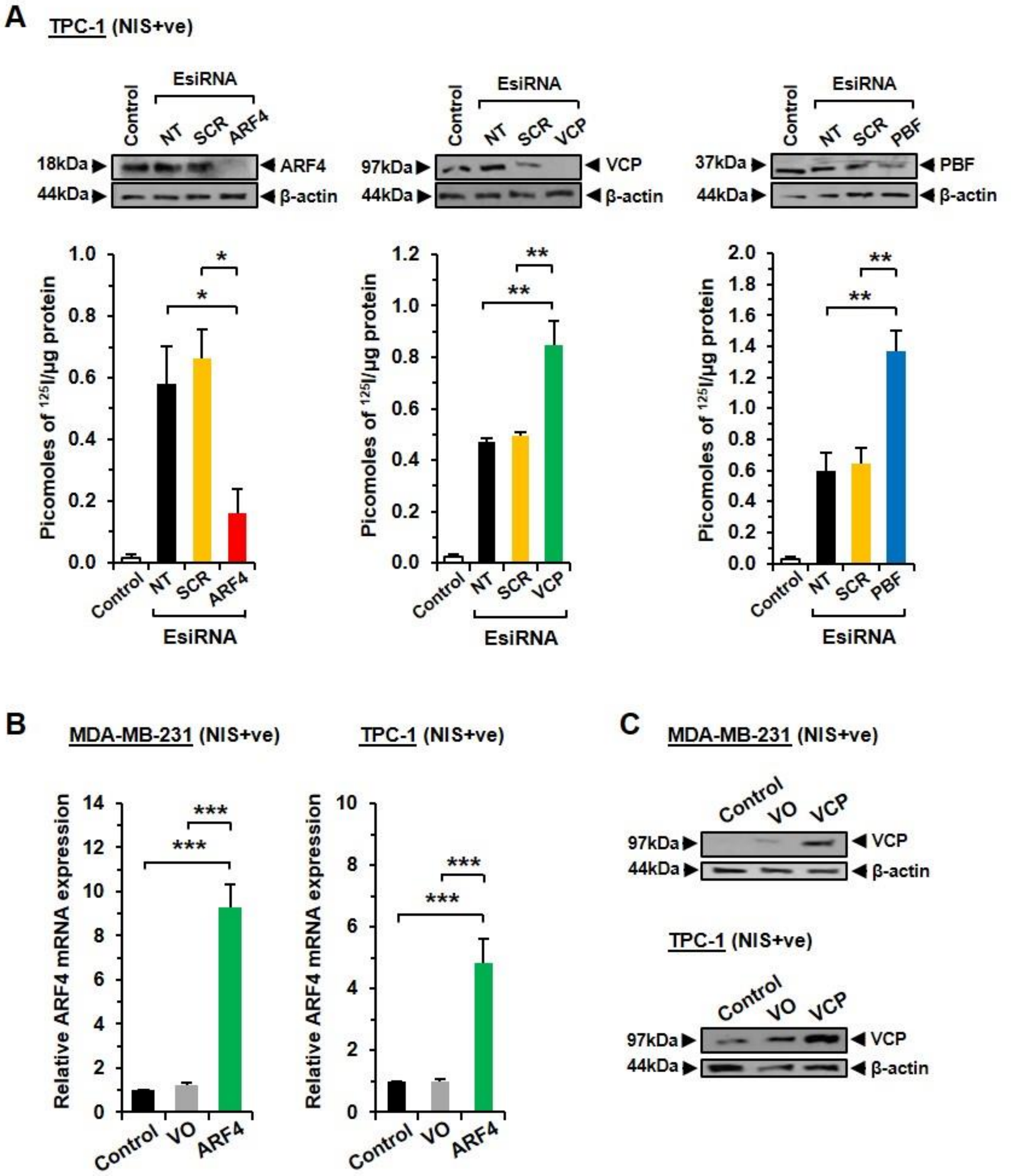


**Figure S2. siRNA ablation of ARF4 and VCP alters $^{125}$I uptake in thyroid cells.** (**A**) RAI uptake and Western blot analysis of TPC-1 (NIS+ve) cells transfected with esiRNA specific for ARF4, VCP or PBF alongside Scr siRNA and non-transfected (NT) TPC-1 (NIS+ve) cells. Control - TPC-1 (NIS-ve) cells. *, $P < 0.05$; **, $P < 0.01$, ANOVA with post hoc analysis. (**B**) Real-time PCR to determine *ARF4* mRNA expression in cells transfected with ARF4 or VO for 48 hours. Control as in (**A**). ***, $P < 0.001$. (**C**) Western blot analysis of VCP protein levels of cells transfected with VCP or VO for 48 hours. Control as in (**A**). All data presented as mean ± SEM from three independent experiments.

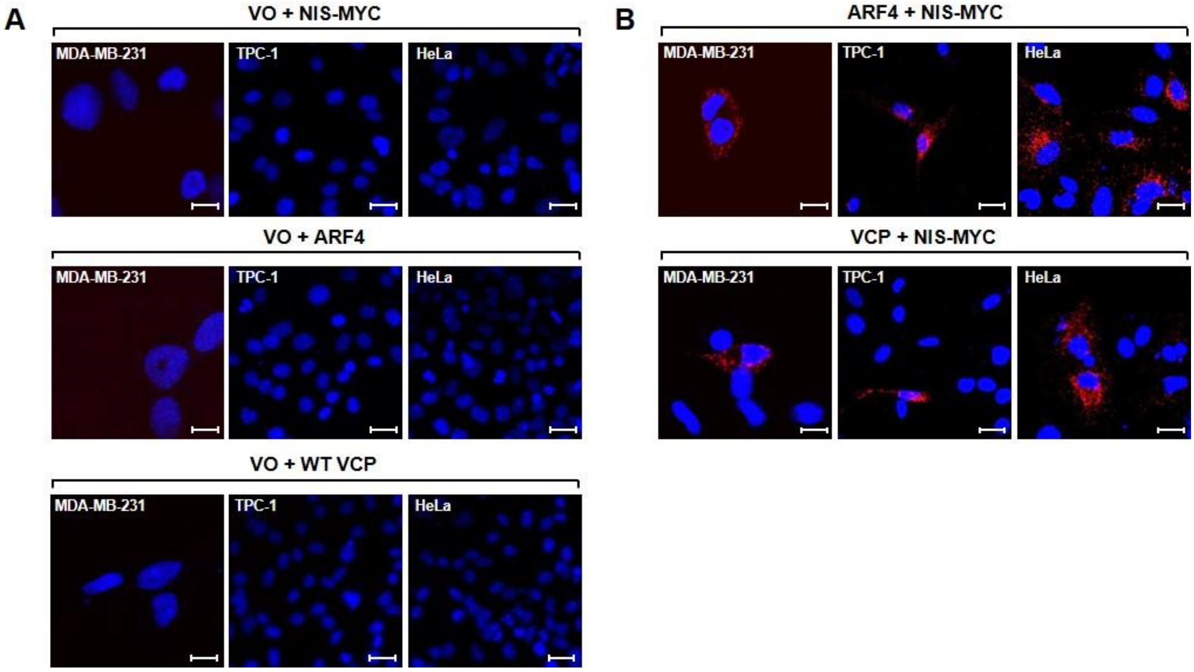


**Figure S3. Novel functional partners ARF4 and VCP specifically interact with NIS.** (**A**) Control PLA assays showing a lack of interaction between VO with NIS-MYC (upper), ARF4 (middle) or VCP (lower) in MDA-MB-231, TPC-1 and HeLa cell lines at 48 hours post-transfection. Blue indicates DAPI nuclear staining. Magnification, 100X. Scale bars, 10 μM. (**B**) PLA assays demonstrating specific interaction between NIS-MYC and ARF4 (upper) or VCP (lower) at 48 hours post-transfection in MDA-MB-231, TPC-1 and HeLa cell lines. Red fluorescent spots indicate specific interactions. Blue indicates DAPI nuclear staining. Magnification, 100X. Scale bars, 10 μM.

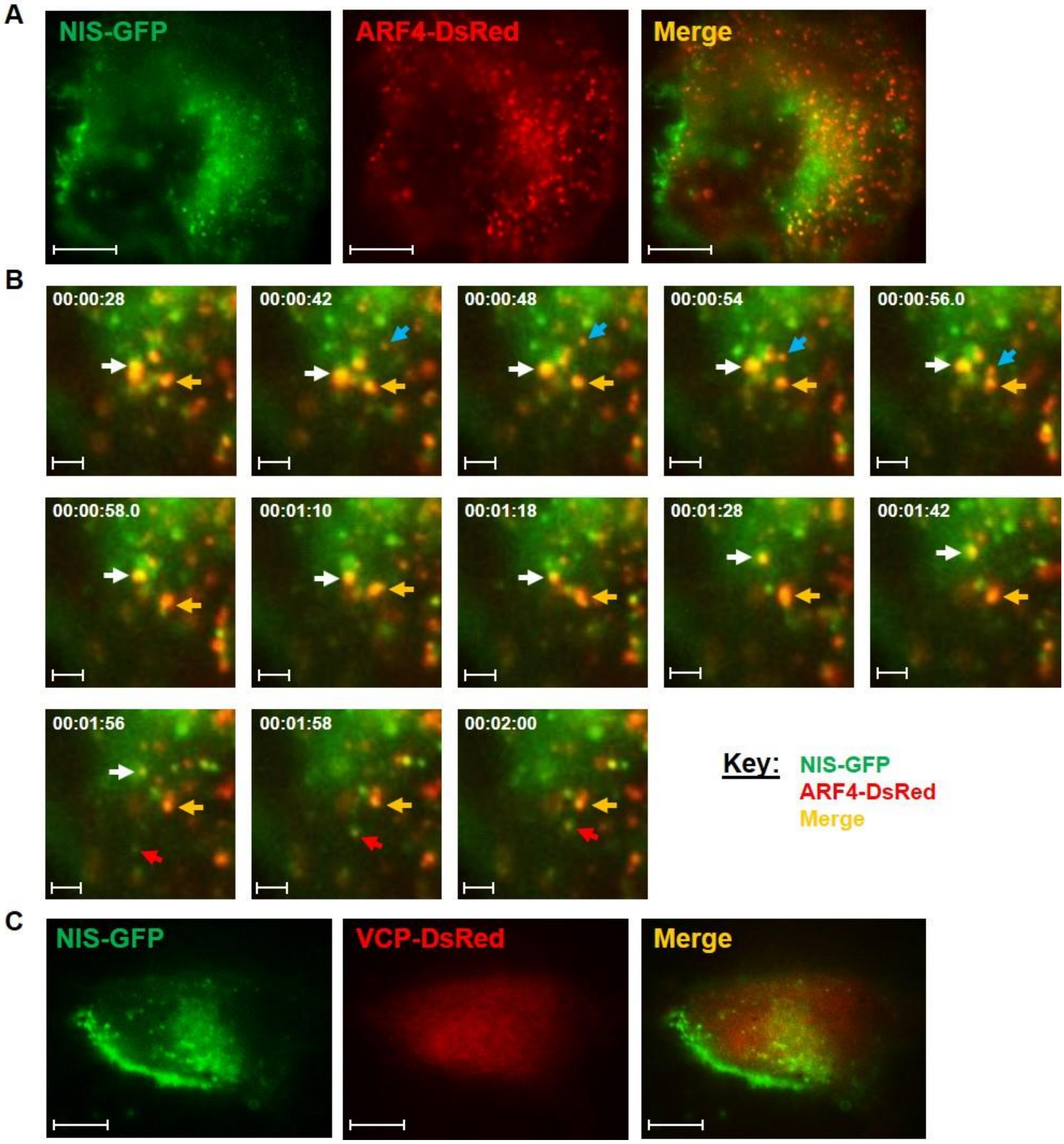


**Figure S4. NIS traffics with ARF4, but not VCP, at the PM.** (**A**) Representative image of a whole HeLa cell using HiLo microscopy demonstrating vesicles at the PM after 48 hours post-transfection of NIS-GFP (green) and ARF4-dsRED (red) with co-localisation in yellow. Scale bars, 10 μM. (**B**) Chronological magnified still frames of PM regions obtained from live cell HiLo microscopy at video capture times indicated (hr:min:sec) following co-transfection in HeLa cells as described in (**A**). Arrowheads indicate co-incident trafficking of vesicles containing NIS-GFP (green) and ARF4-dsRED (red) at the PM, demonstrated by co-localisation (yellow). Scale bars, 10 μM. (**C**) Representative image of a whole HeLa cell using HiLo microscopy following co-transfection of NIS-GFP (green) and VCP-dsRED (red). Scale bars, 10 μM.

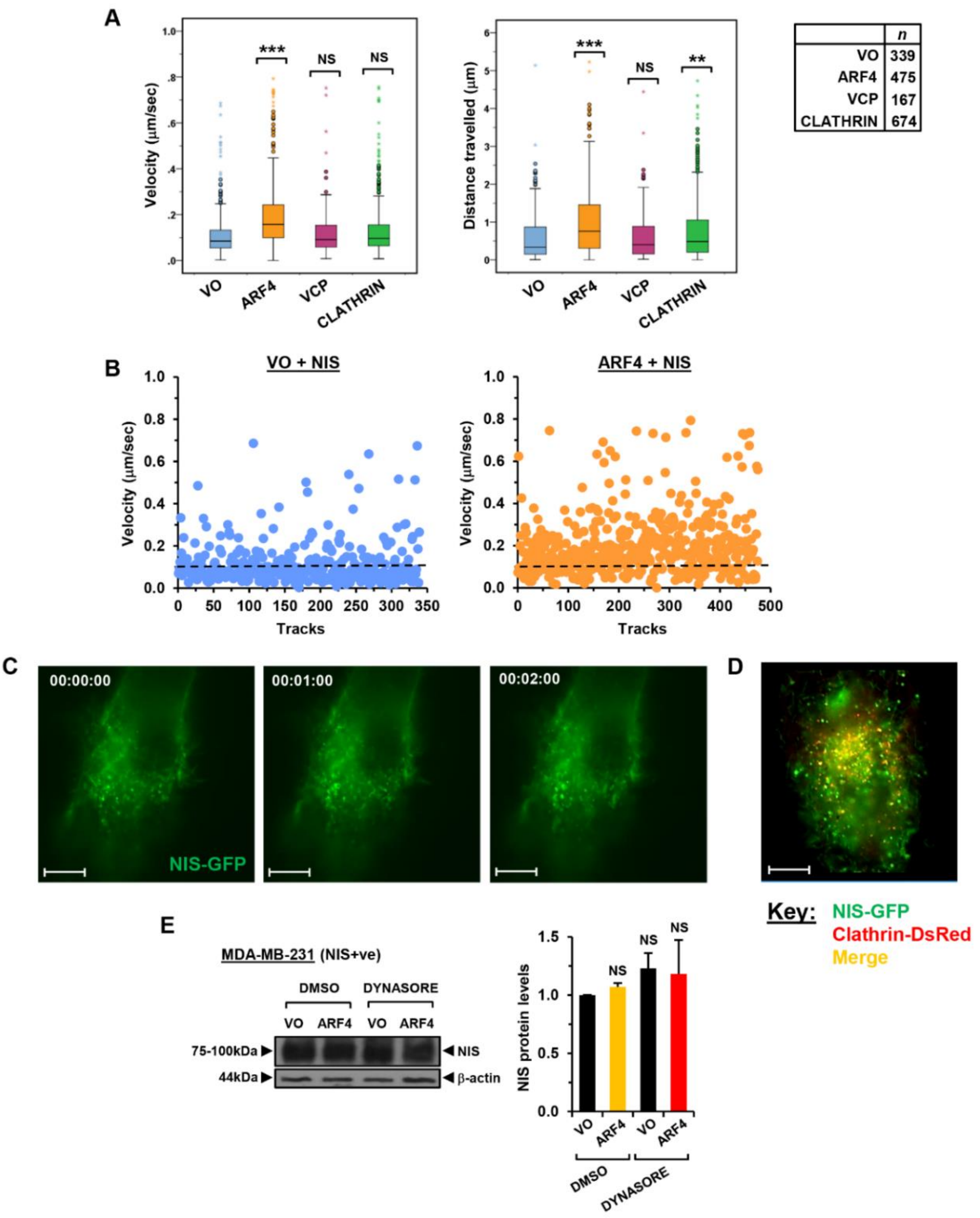


**Figure S5. ARF4 increases the velocity and distance travelled of NIS-GFP positive vesicles.** (**A**) Box and whisker plots of velocity (µm/sec) and distance travelled (µm) of NIS vesicles following co-transfection of HeLa cells with NIS-GFP and ARF4-dsRED (*n* = 475), VCP-dsRED (*n* = 167), Clathrin-dsRED (*n* = 674) or VO (*n* = 339). NS, not significant; **, $P < 0.01$; ***, $P < 0.001$. (**B**) Dot plots of velocity (µm/sec) of individual NIS vesicles co-transfected with ARF4-dsRED (*n* = 475) or VO (*n* = 339) in HeLa cells, as outlined in (**A**). (**C**) Representative images of a whole cell at video capture times indicated (hr:min:sec) obtained using HiLo microscopy. NIS vesicles at the PM following co-transfection of NIS-GFP (green) and VO in HeLa cells. Scale bars, 10 µm. (**D**) Representative image of the PM of a HeLa cell obtained by HiLo microscopy following co-transfection with NIS-GFP (green) and Clathrin-dsRED (red), with co-localisation in yellow. Scale bars, 10 µm. (**E**) Western blot analysis of NIS expression levels (left) quantified by densitometry (right) in TPC-1 (NIS+ve) cells transfected with ARF4 or VO prior to treatment with Dynasore or DMSO for 1 hour. NS, not significant. Data presented as mean ± SEM from three independent experiments.

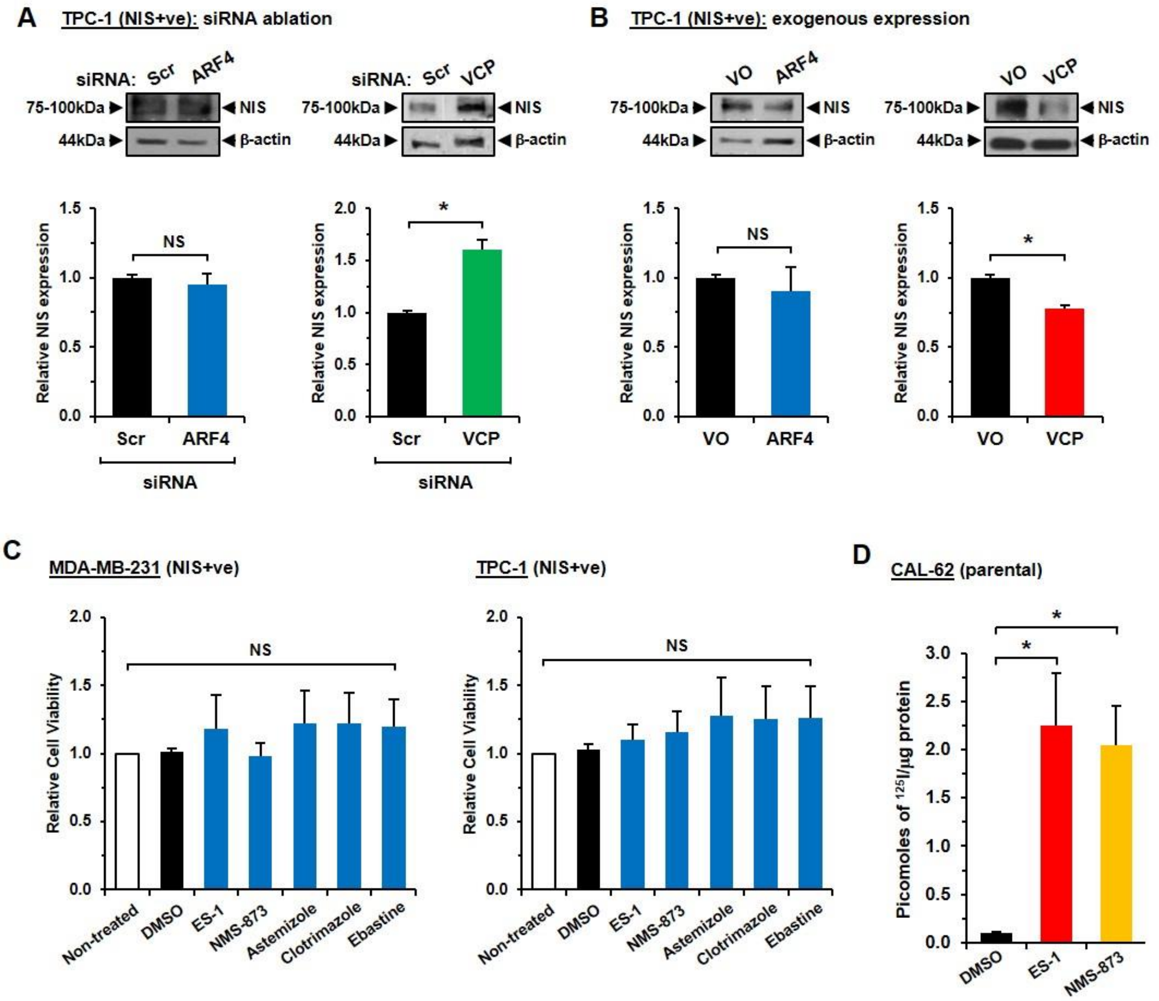


**Figure S6. ARF4 and VCP differentially affect NIS expression, but VCP inhibitors do not alter cell viability.** (**A**) Upper, Western blot analysis of NIS expression in TPC-1 (NIS+ve) cells transfected with ARF4 (left) and VCP (right) siRNA or Scr siRNA. Lower, densitometry of three representative Western blots. NS, not significant; *, $P < 0.05$, Student's t-test. (**B**) Same as (**A**), but cells transfected with ARF4 (left) and VCP (right) or VO. (**C**) CellTiter-Glo® luminescent cell viability assay showing no significant reduction in TPC-1 (NIS+ve) cell viability following treatment with 2.5 µM ES-1, 5 µM NMS-873, 0.25 µM Astemizole, 0.25 µM Clotrimazole, 0.5 µM Ebastine or DMSO for 24 hours. Non-treated control received no drug treatment. NS, not significant, ANOVA with post hoc analysis. (**D**) RAI uptake in parental CAL-62 cells treated with DMSO, 2.5 µM ES-1 or 5 µM NMS-873 for 24 hours prior to addition of $^{125}$I. *, $P < 0.05$, ANOVA with post hoc analysis. All data presented as mean ± SEM from three independent experiments.

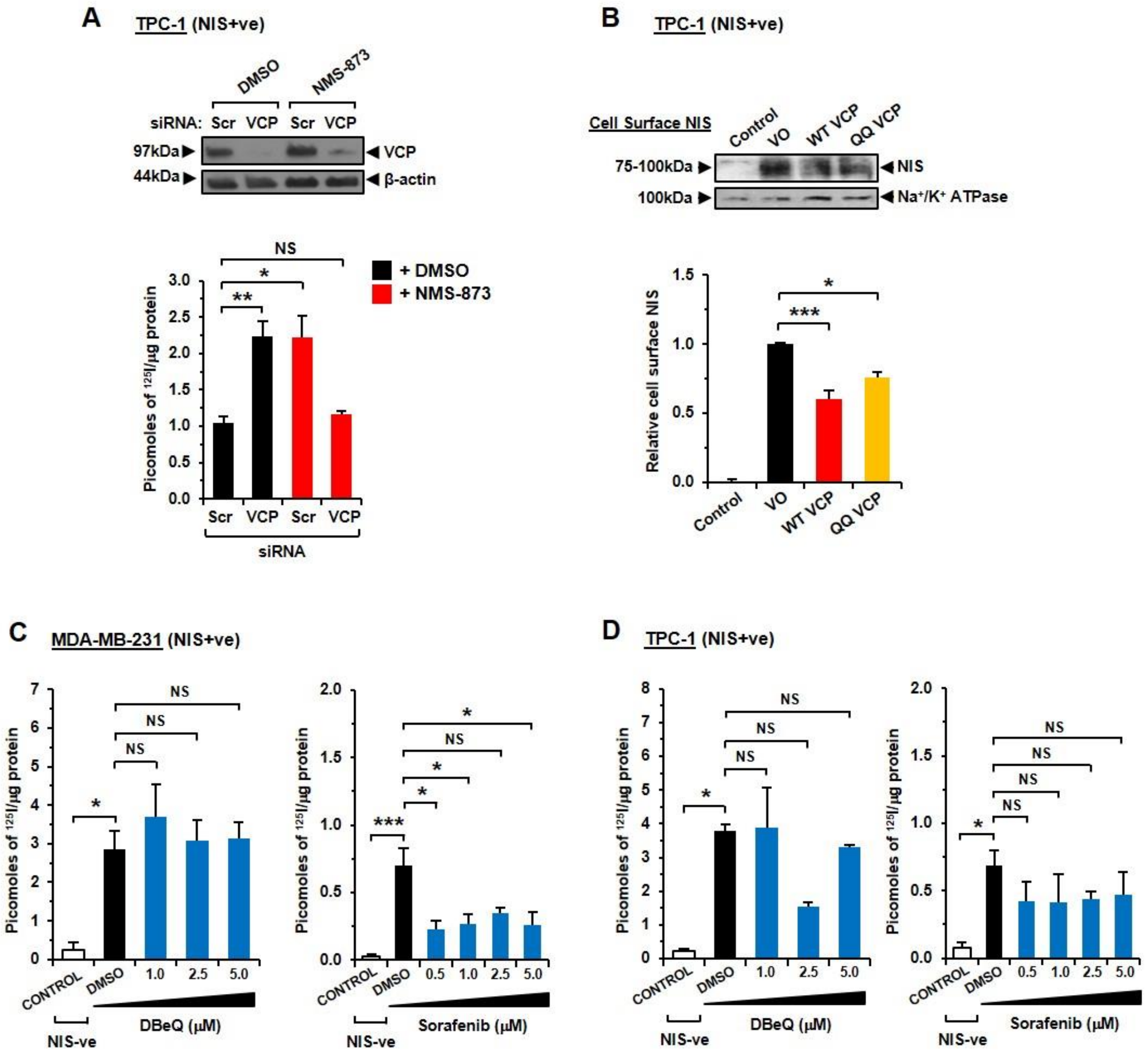


**Figure S7. VCP inhibitors require VCP expression to exert their effect on NIS function.** (**A**) RAI uptake (lower) in TPC-1 (NIS+ve) cells transfected with VCP or Scr siRNA for 48 hours prior to treatment with 5 μM NMS-873 for 24 hours. Western blot analysis (upper) of VCP expression following transfection and drug treatment as above, to confirm VCP depletion following transfection with VCP siRNA. NS, not significant; *, $P < 0.05$; **, $P < 0.01$, ANOVA with post hoc analysis. (**B**) CSBA analysis by Western blot (upper) to determine PM NIS expression in TPC-1 (NIS+ve) cells following transfection with wild type VCP (WT VCP), ATPase-deficient dominant-negative VCP mutant (QQ VCP) or VO. Control - TPC-1 (NIS-ve) cells. Quantification of three independent Western blots using densitometry to assess cell surface NIS expression relative to $Na^+/K^+$ ATPase (lower). Data presented as mean NIS levels ± SEM. *, $P < 0.05$; ***, $P < 0.001$. (**C**) MDA-MB-231 (NIS+ve) cells treated with a range of doses of DBeQ (μM) for 24 hours prior to addition of $^{125}$I. Control – MDA-MB-231 (NIS-ve) cells. NS, not significant; *, $P < 0.05$; ***, $P < 0.001$, ANOVA with post hoc analysis. (**D**) Same as (**C**), but TPC-1 (NIS+ve) cells. All data presented as mean ± SEM from three independent experiments.

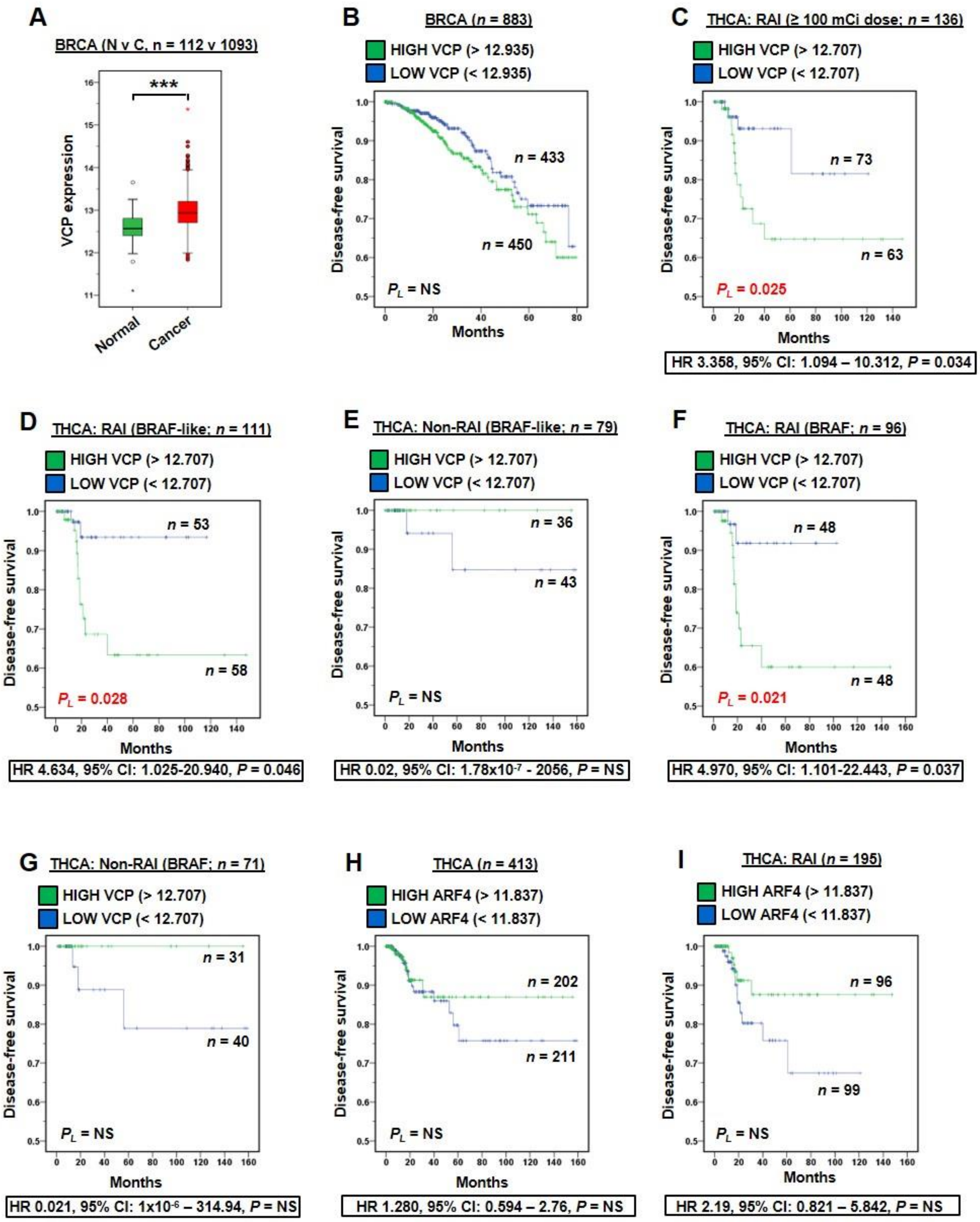


**Figure S8. VCP expression correlates with worse DFS in PTC with BRAF alterations.** (**A**) Box whisker plots of VCP expression in normal (*n* = 112) versus cancer (*n* = 1093) in BRCA TCGA data. (**B**) Kaplan-Meier plot of DFS for BRCA with high (Q3Q4) versus low (Q1Q2) VCP expression for the entire BRCA cohort. NS, not significant. (**C-G**) DFS for THCA with high (Q3Q4) versus low (Q1Q2) VCP expression for RAI-treated patients (≥100 mCi) (**C**), RAI-treated patients with a BRAF-like genetic signature (**D**), non-RAI treated patients with a BRAF-like genetic signature (**E**), RAI-treated patients with BRAF mutation (**F**) and non-RAI treated patients with a BRAF mutation (**G**). Hazard ratios (HR) ± 95% CI are indicated below each panel. NS, non-significant. (**H-I**) DFS for THCA with high (Q3Q4) versus low (Q1Q2) ARF4 expression for the entire PTC cohort (**H**) and RAI-treated patients (**I**). Hazard ratios (HR) ± 95% CI are indicated below each panel. NS, non-significant.

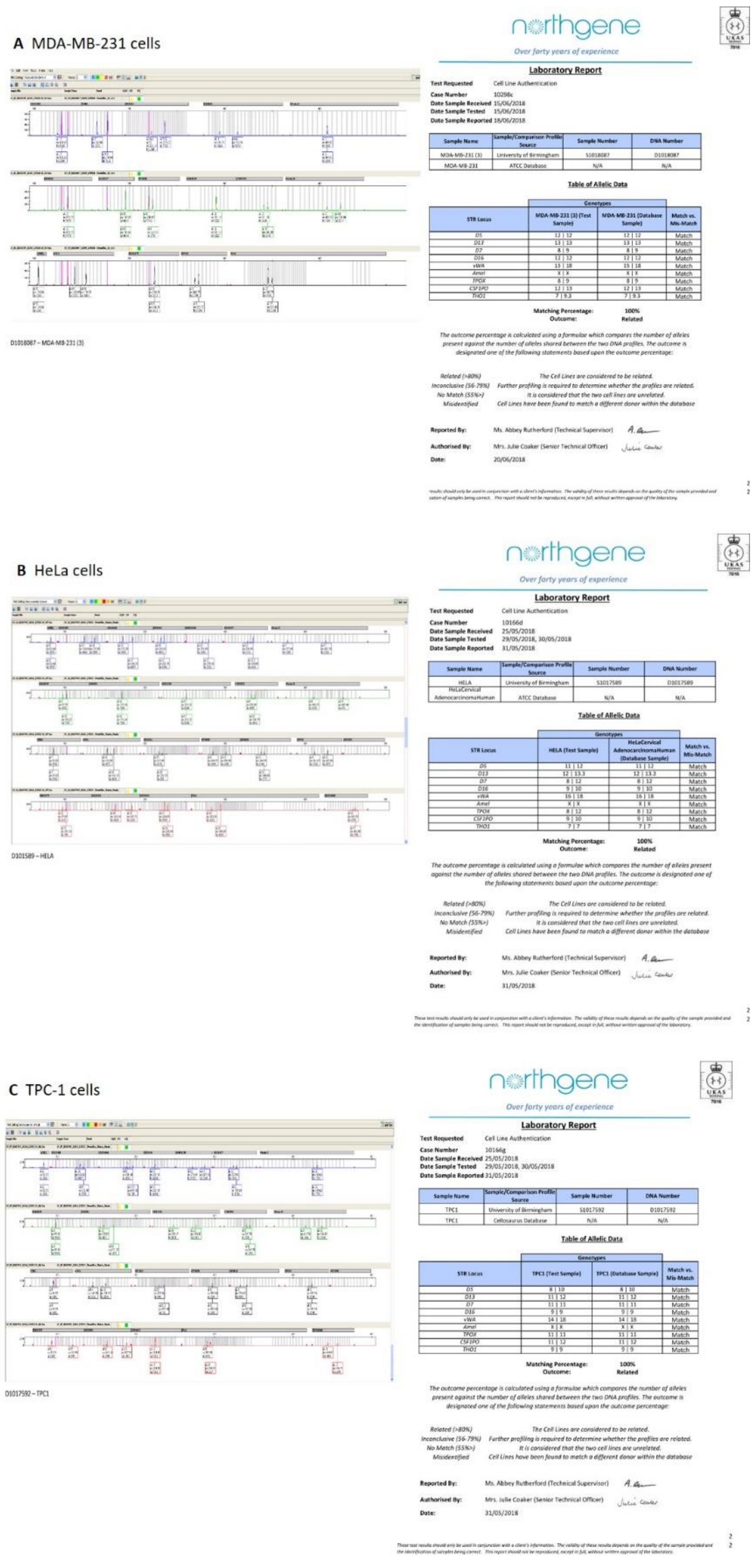


**Figure S9. Short tandem repeat (STR) profiling of cell lines used in study.** Panels **A** to **C** show a representative copy of the electropherogram and cell line DNA typing report obtained following Short Tandem Repeat (STR) analysis by the accredited company NorthGene (Newcastle upon Tyne). Authenticity was assessed by comparing the generated STR profile with the source STR profiles present in the American Type Culture Collection (ATCC), Cellosaurus and the Deutsche Sammlung von Mikroorganismen und Zellkulturen (DSMZ).

#### Supplementary tables

| Gene | Protein | Whole cell (n=3) | | | Plasma membrane (n=3) | | | Combined (n=6) | | |
|---|---|---|---|---|---|---|---|---|---|---|
| | | Score[A] | Peptides[B] | SC (%)[C] | Score[A] | Peptides[B] | SC (%)[C] | Score[A] | Peptides[B] | SC (%)[C] |
| SLC5A5 | Sodium/iodide symporter | 458.9 | 25 | 12.1 | 110.3 | 7 | 4.5 | 284.6 | 32 | 8.3 |
| ARF4[D] | ADF-ribosylation factor 4 | 62.3 | 5 | 10 | 120.9 | 8 | 15.4 | 91.6 | 13 | 12.7 |
| VCP[E] | Valosin-containing protein | 161.0 | 12 | 5.6 | - | - | - | 80.6 | 12 | 2.8 |
| EHD1[F] | EH domain-containing protein 1 | - | - | - | 110.9 | 8 | 6.1 | 55.4 | 8 | 3.1 |
| RHEB | GTP-binding protein Rheb | 6.9 | 1 | 2.9 | 114.9 | 6 | 13.7 | 60.9 | 7 | 8.3 |
| PDCD6IP | Programmed cell death 6-interacting protein | - | - | - | 105.7 | 7 | 3.1 | 52.9 | 7 | 1.6 |
| ANXA1 | Annexin A1 | - | - | - | 105.1 | 7 | 10.1 | 52.6 | 7 | 5.1 |
| SAR1A | GTP-binding protein SAR1a | - | - | - | 102.8 | 7 | 14.9 | 51.4 | 7 | 7.5 |
| PP1G | Serine/threonine-protein phosphatase PP1-gamma catalytic subunit | - | - | - | 61.4 | 7 | 9.6 | 30.7 | 7 | 4.8 |
| LDHA | L-lactate dehydrogenase A chain | 136.2 | 6 | 6.9 | - | - | - | 68.1 | 6 | 3.5 |
| PPIB | Peptidyl-prolyl cis-trans isomerase B | - | - | - | 126.8 | 6 | 12.2 | 63.4 | 6 | 6.1 |
| COPB1 | Coatomer subunit beta | - | - | - | 105.1 | 6 | 2.0 | 52.6 | 6 | 1.0 |
| GNB2L1[G] | Guanine nucleotide-binding protein subunit beta-2-like 1 | 44.2 | 2 | 2.8 | 59.8 | 4 | 5.3 | 52.0 | 6 | 4.1 |
| BAG2 | BAG family molecular chaperone regulator 2 | 70.8 | 5 | 7.4 | 30.0 | 1 | 1.7 | 50.4 | 6 | 4.6 |
| NT5E[H] | 5'-nucleotidase ecto | - | - | - | 84.6 | 6 | 4.3 | 42.3 | 6 | 2.2 |

**Table S1. Top hits for putative NIS interactors identified by mass spectrometry.** [A]Score: Mascot score assesses the likelihood that the identified peptide match with the in silico predicted peptide is significant. Mean scores for whole cell lysates (*n* =3), plasma membrane proteins (*n* = 3) and the combination of these (*n* = 6) are shown. [B]Peptides: number of peptides of a protein identified within the MS/MS screen, which is dependent on multiple factors, including the amount of a given protein within a sample, trypsin cleavage efficiency of the protein as well as observation and identification of the subsequent peptides by MS/MS. [C]SC(%): Sequence coverage (SC) is a measure of how much the theoretical protein's amino acid sequence is covered by the experimental peptides identified. [D]ARF4: also known as ARF2. [E]VCP: also known as P97, TERA and CDC48. [F]EHD1: also known as PAST1. [G]GNB2L1: also known as RACK1. [H]NT5E: also known as 5-NT, NT5, NTE and CD73.

| Target | Type of siRNA | Company | ID |
|---|---|---|---|
| ARF4 | MISSION esiRNA[A] | Sigma-Aldrich | EHU038521 |
| VCP | MISSION esiRNA | Sigma-Aldrich | EHU071991 |
| EHD1 | MISSION esiRNA | Sigma-Aldrich | EHU093311 |
| NT5E | MISSION esiRNA | Sigma-Aldrich | EHU023471 |
| GNB2L1 | MISSION esiRNA | Sigma-Aldrich | EHU102911 |
| ARF4 | ONTARGETplus siRNA[B] | Dharmacon | L-011582-00-0005 |
| VCP | MISSION siRNA (pooled)[C] | Sigma-Aldrich | Hs01_00118726; Hs02_00343009; Hs01_00118728 |
| PBF | Silencer siRNA (pooled)[D] | ThermoFisher Scientific | s2233; s2234 |
| Scrambled | Negative control siRNA | ThermoFisher Scientific | AM4635 |

**Table S2. siRNAs used in study.** [A]MISSION endo-ribonuclease prepared siRNA (esiRNA) are a heterogeneous mixture of siRNAs that all target the same mRNA sequence. MISSION esiRNA were used in an initial screen to investigate the impact on NIS function of five putative interactors identified by MS/MS in MDA-MB-231 NIS+ve cells (Figure 1, A-D). [B]A pre-prepared mixture of four siRNAs provided as a single reagent were used to confirm that ARF4 ablation decreased radioiodine uptake in TPC-1 and MDA-MB-231 NIS+ve cells as well as primary human thyrocytes (Figure 1E). [C]Pooled reagent of three different MISSION siRNAs targeted to VCP. Designed using algorithm from Rosetta Inpharmatics. Approximate positioning where siRNA targets gene are at nucleotides 768 (Hs01_00118726), 1908 (Hs02_00343009) and 2548 (Hs01_00118728). Used to confirm that VCP ablation increased radioiodine uptake in NIS+ve cell types and primary human thyroid cells (Figure 1G). [D]Pooled reagent of two different silencer siRNAs targeted to PBF. Previously, we demonstrated efficient silencing of PBF mRNA (> 90%) using this siRNA combination in cells (*81*).

| Antibody | Clone | Supplier |
|---|---|---|
| PBF[A] | Rabbit polyclonal; Custom | Eurogentec |
| NIS[A,B] | Rabbit polyclonal [24324-1-AP] | ProteinTech |
| ARF4[A,C] | Rabbit polyclonal [11673-1-AP] | ProteinTech |
| VCP[A,C] | Rabbit polyclonal [10736-1-AP] | ProteinTech |
| HA[A] | Mouse monoclonal [16B12] | BioLegend |
| MYC[C] | Mouse monoclonal [9B11] | Cell Signaling Technology |
| $Na^+/K^+$ ATPase[A,D] | Rabbit monoclonal [EP1845Y] | Abcam |
| α-tubulin[A] | Mouse monoclonal [B7] | Santa Cruz Biotechnology |
| β-actin[A] | Mouse monoclonal [AC-15] | Sigma-Aldrich |

**Table S3. Primary antibodies used in study.** [A]Primary antibodies used in Western blotting. To quantify detected bands by densitometry, blots were scanned, keeping all scanning parameters the same, and analyzed using ImageJ software. [B]Anti-NIS antibody used to co-immunoprecipitate NIS interactors prior to identification by MS/MS. [C]Antibodies used to detect specific NIS interactions in proximity-ligation assays. [D]Anti-$Na^+/K^+$ ATPase antibody used as a plasma membrane loading control.

| Vector[1] | Gene | Source | Notes |
|---|---|---|---|
| pcDNA3.1(+) | NIS-HA | Previously described (Smith et al., 2013) | |
| pcDNA3.1(+) | AAAK NIS-HA[A] | This study | $_{574}$VAPK$_{577}$ motif mutated to $_{574}$AAAK$_{577}$ |
| pcDNA3.1(+) | ALAS NIS-HA[A] | This study | $_{475}$VLPS$_{478}$ motif mutated to $_{475}$ALAS$_{478}$ |
| pcDNA3.1(+) | NIS-MYC | Previously described (Smith et al., 2013) | |
| pEGFP-N3 | NIS | Dr Takahiko Kogai (Dokkyo Medical University) | |
| pCMV6-XL5 | ARF4 | #SC119092; Origene | |
| pDsRed-Express-N1 | ARF4[B] | This study | BamHI-HindIII restriction |
| pcDNA3.1(+) | VCP (rat)[C,D] | Dr Yihong Ye (National Institutes of Health) | |
| pcDNA3.1(+) | QQ VCP (rat)[C,E] | Dr Yihong Ye (National Institutes of Health) | |
| pcDNA3.1(+) | WT VCP (human)[F] | This study | N794S substitution |
| pcDNA3.1(+) | QQ VCP (human)[E,F] | This study | N794S substitution |
| pEGFP-N1 | VCP | #23971; Addgene | |
| pDsRed-Express-N1 | VCP[B] | This study | BamHI-HindIII restriction |
| pDsRed-Express-N1 | Clathrin[G] | Dr Natalie Poulter (University of Birmingham) | |

**Table S4. Nucleic acids used in study.** Expression vectors without an inserted gene of interest were used as vector only (VO) controls in study. [A]AAAK and ALAS mutants of NIS-HA were generated using the QuikChange Site-directed Mutagenesis Kit (Agilent Technologies). Mutagenesis primers used: $_{574}$AAAK$_{577}$ NIS-HA - 5' GCACGGCAGACAGCATCAGCGGCCGCCAAGGAAGAAGTGGC 3' (forward) and 5' GCCACTTCTTCCTTGGCGGCCGCTGATGCTGTCTGCCGTGC 3' (reverse); $_{475}$ALAS$_{478}$ NIS-HA - 5' CAT GAG GGC CCT GGC ATC GTC G 3' (forward) and 5' CGA CGATGCCAGGGCCCTCAT G 3' (reverse). [B]ARF4 and VCP cDNAs were subcloned into the pDsRed-Express-N1 vector for use in HiLo microscopy. VCP cDNA was excised from the VCP(wt)-EGFP plasmid (#23971; Addgene) and ligated into pDsRed-Express-N1 by BamHI-HindIII restriction, whereas ARF4 cDNA was PCR amplified from the ARF4-pCMV6-XL5 plasmid (#SC119092; Origene) prior to ligation into pDsRed-Express-N1 by BamHI-HindIII restriction. PCR primers used: 5' GCCAAGCTTCCACCATGGGCCTCACTATCT C 3' (forward) and 5' CGGGGATCCTCATTAACGTTTTGAAAGCTC 3' (reverse). [C]All plasmids code for the full wild type amino acid sequence for the human gene of interest, unless otherwise stated. [D]For further information see references (*82*-*83*). [E]The QQ VCP mutant is defective in ATP-hydrolysis due to two point mutations within the ATPase domains at E305Q and E578Q within the Walker B motifs (*82*). [F]Rat and human VCP have >99.8% amino acid similarity with only one mismatch at site 794 (a serine in the human sequence and asparagine in the rat sequence). Human WT VCP and QQ VCP cDNA were generated from rat WT VCP and QQ VCP sequences by site-directed mutagenesis. Primers used: 5' CACAGGTGGCAGTGTGTACAC 3' (forward) and 5' GTGTACACACTGCCACCTGTG 3' (reverse). [G]For further information see reference (*84*).

#### Supplementary movies (attached separately)

**Movie S1:** Significant co-localisation and trafficking of ARF4-dsRED and NIS-GFP proteins in co-incident vesicles at the plasma membrane in HeLa cells

**Movie S2:** Lack of co-trafficking for VCP-dsRED and NIS-GFP suggests that the site of functional interaction between VCP and NIS is distant to the plasma membrane in HeLa cells

**Movie S3:** The sodium iodide symporter NIS is endosomally trafficked in association with clathrin
